# Cytogenetic constraints on hybridization: A meta-analysis investigating the role of chromosome number in monocot hybrid evolution using a newly developed tool, the ploidy deviation index (PDI)

**DOI:** 10.1101/2025.11.13.688279

**Authors:** Justin Scholten, Arianna Sprenger, Ava Perez, Olivia Hullihen, Chelsea D. Specht

## Abstract

**Background and aims:** Hybridization is a major driver of plant diversity, yet the role of cytogenetic compatibility, particularly differences in chromosome number, remains poorly understood. Differences in parental chromosome number can present barriers to hybrid formation by disrupting meiotic stability, but the extent to which biological and ecological factors influence the chromosomal architecture of hybrids remains poorly quantified, especially in monocots. This study aims to investigate how chromosome number divergence interacts with biological and ecological factors to shape hybrid formation in monocots, using a novel quantitative metric, the Ploidy Deviation Index (PDI), to standardize comparisons of hybrid cytogenetic architecture.

**Material and methods:** We developed and applied the PDI, a continuous index quantifying chromosome-number deviation between a hybrid and its two parents, across approximately 200 hybrid cases with documented parental karyotypes. Hybrids were categorized as homoploid, uniparentally homoploid, intermediate, or polyploid based on their PDI values. We analyzed the distribution of PDI scores in relation to type of hybrid origin (natural vs. artificial), growth habit, size of the genus (a proxy for richness), and range of chromosome number within a genus (proxy for diversity). Comparisons across categories employed Anderson–Darling k-sample tests, multinomial logistic regression, and Mann–Whitney U tests to determine significance.

**Key results:** Homoploid hybrids were found to be the most frequent. We found no significant difference in PDI distributions between natural and artificial hybrids. Significant variation in PDI distributions was found among growth habits, with aquatic hybrids more likely to be homoploid and geophytic hybrids showing higher proportions of polyploidy. Intermediate hybrids were common in larger genera with broader chromosome-number ranges, whereas polyploid hybrids showed the highest PDI values in large and karyotypically diverse genera.

**Conclusion:** These results challenge long-held assumptions that polyploidy dominates hybrid formation and reveal that homoploid and intermediate chromosomal configurations are common in monocots. The PDI framework offers a powerful, standardized approach for assessing cytogenetic constraints on hybridization, with implications for systematics, evolutionary biology, and conservation.

## INTRODUCTION

Hybridization has long been recognized as a major evolutionary force in plants, with the potential to drive speciation, facilitate adaptation, and generate phenotypic novelty (Anderson 1949; Rieseberg 1997; Abbott et al. 2013). In monocots—one of the two major clades of angiosperms harboring 30% of flowering plant diversity—hybridization has been shown to play a particularly significant role in horticultural and natural systems alike, yielding a diverse array of hybrid taxa that span the range of reproductive compatibility, genome structure, and ecological niche utilization (Soltis and Soltis, 2009; Pfennig et al. 2016). Yet while numerous studies have documented the frequency and outcomes of hybridization events, the role of cytogenetic compatibility—especially differences in chromosome number—remains underexplored at a broad scale. Chromosome number variation can represent a significant postzygotic barrier to successful hybrid formation and stabilization, yet its constraints, tolerances, and consequences for hybrid lineages remain poorly quantified across monocots.

Cytogenetic compatibility between parental species is widely recognized as a prerequisite for stable meiosis in hybrids (Grant, 1981; Ramsey and Schemske, 1998). Hybrids between species with different chromosome numbers often suffer from reduced fertility, chromosomal missegregation, and unbalanced gametes (Stebbins, 1950; Levin, 2002). However, the extent to which chromosome number divergence constrains hybridization in practice remains uncertain, especially across the structural diversity and evolutionary breadth of monocotyledonous plants. Monocots include some of the most cytogenetically variable lineages, from genera like *Allium* L., *Carex* L., and *Orchis* Tourn. ex L., where extensive dysploidy, polyploidy, and chromosome fusion/fission events are common (Peruzzi and Eroğlu, 2013; Márquez-Corro et al., 2019). These structural rearrangements may either inhibit hybridization or be tolerated under particular genomic or ecological contexts. Whether patterns of natural and artificial hybridization reflect strict cytogenetic constraints or greater tolerance for karyotypic disparity remains an open question.

Despite its potential importance, no meta-analysis has yet systematically quantified the relationship between chromosome number and hybrid formation in monocots. Past reviews have focused on the prevalence of polyploidy in plant evolution more broadly (Soltis et al., 2004; Wood et al., 2009), but few have distinguished among distinct cytogenetic scenarios—such as homoploid hybridization, polyploid hybrid formation, or intermediate and uniparentally homoploid systems. Moreover, the distinction between natural and artificial (particularly horticultural) hybrids is rarely made, though this distinction offers important insights into what cytogenetic configurations are biologically possible versus what is only realized under human intervention. This study represents the first attempt to quantitatively assess the role of chromosome number disparity in monocot hybrid formation across hundreds of reported cases.

Importantly, this study introduces a novel mathematical framework for quantifying chromosome number disparity between hybrids and their parental species—a “ploidy deviation index” (PDI). While previous studies have occasionally noted cytogenetic differences informally or through basic ratios (Ramsey and Schemske, 2002), this is the first effort to formalize a continuous metric that can distinguish among homoploid, polyploid, and intermediate cases on a standardized scale. This framework not only captures the direction and magnitude of chromosome number shifts but can be applied to any hybrid system where hybrid and parental karyotypes are known. By calculating PDI for each hybrid case, we can assess whether particular disparities (i.e. deviation scores) are overrepresented in specific life histories, thereby shedding light on the tolerance or rejection of cytogenetic divergence in hybrid formation.

The evolutionary implications of such patterns are profound. Homoploid hybridization, though historically believed to be rare, has been implicated in ecological novelty and the generation of new species without the reproductive isolation afforded by polyploidy (Gross and Rieseberg, 2005; Feliner et al., 2017). Polyploid hybridization, conversely, often circumvents sterility by doubling chromosome sets and may play a disproportionate role in long-term hybrid lineage stabilization (Stebbins, 1971; Soltis and Soltis, 1999). Uniparental homoploidy and intermediate karyotypes represent more ambiguous configurations that may offer insights into meiotic tolerance, structural homology, or cryptic genomic flexibility—features not typically captured in binary models of hybrid fertility or success.

Understanding cytogenetic constraints also has direct implications for systematics, conservation, and horticulture. Misidentified hybrids or taxa of uncertain status often reside in taxonomic limbo due to overlooked chromosomal mismatches (Weiss-Schneeweiss and Schneeweiss, 2013), and the ability to predict hybrid viability based on karyotype compatibility could aid in delimiting species boundaries, identifying unique cytotypes, and managing hybrid zones. In horticulture, chromosome number has long informed breeding programs (e.g., in *Lilium* Tourn. ex L., Orchidaceae Juss., and *Tulipa* L.), but the extent to which artificial hybrids deviate from natural cytogenetic parental species is not well documented. This analysis offers a broad-scale empirical foundation to evaluate the role of chromosome number in shaping the landscape of viable hybridization, impacting efforts in cultivation and conservation.

By synthesizing cytogenetic data from hundreds of documented hybrid systems, this study provides the first comprehensive meta-analysis of chromosomal compatibility in monocot hybrid evolution. It also proposes a new tool for assessing hybrid cytogenetic architecture, facilitating future studies that aim to quantify the evolutionary, developmental, and practical implications of chromosomal divergence. Ultimately, this work aims to move the discussion of hybrid viability beyond anecdotal evidence and toward a more predictive, quantitative understanding of cytogenetic constraints in plant evolution.

## MATERIALS AND METHODS

### Ploidy deviation index (PDI)

To investigate the degree to which chromosomal differences constrain or shape hybrid formation in monocots, we developed a continuous, standardized metric termed the *Ploidy Deviation Index* (PDI). This index quantifies the magnitude and direction of divergence in chromosome number between a hybrid and its two putative parents. Unlike binary classifications of polyploidy or homoploidy, the PDI allows fine-scale comparisons across a continuous gradient of numerical relationships and facilitates meta-analytical treatment of heterogeneous cytogenetic data.

### Mathematical formulation

Let *C_H_* represent the somatic chromosome number of a hybrid taxon, and *C_P1_* and *C_P2_*represent the chromosome numbers of its two parental species. The first step in calculating the PDI is to determine the average squared deviation of the hybrid from each parent:

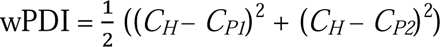

This “weighted PDI” (wPDI) captures the absolute deviation of the hybrid from its cytogenetic parental envelope. Squaring the differences ensures that larger deviations are weighted more heavily—an approach that is biologically justified, as large differences in chromosome number are more likely to cause meiotic irregularities, unbalanced gametes, and reduced fertility (Levin, 2002). In contrast, hybrids with small deviations from both parents may retain partial chromosomal pairing and thus higher potential for reproductive success.

To standardize this value across taxa with different chromosome number scales, the weighted PDI is divided by the squared difference between the parental chromosome numbers, which we denote as *R*, resulting in the normalized PDI (nPDI):

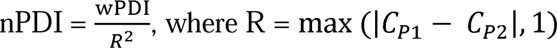

The normalization factor *R* ensures that values are scaled appropriately across hybrid systems with different parental chromosome number ranges. Importantly, it also prevents inflation of the index in lineages with high base numbers and avoids division by zero when *C_P1_* = *C_P2_*, such as in cases of conspecific or closely related homoploid hybrids. Homoploid hybrids, by definition, will always return a score of 0 under this formulation.

### Directional correction

In some cases, hybrids match the chromosome number of one parent exactly but not the other. These “uniparentally homoploid” hybrids may result from successful backcrossing, partial genome elimination, or asymmetric genomic stabilization. To capture this directional similarity, the index is made negative whenever the hybrid matches the chromosome number of one parent exactly, resulting in the sign-corrected PDI (sPDI):

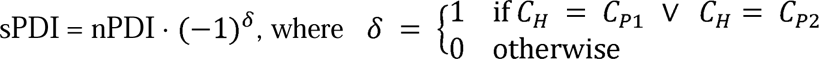

Assigning negative scores to these cases is important for two reasons. First, it reflects the underlying biological asymmetry often observed in such hybrids, where meiotic compatibility or genome dominance favors one parental lineage over the other (Leitch and Leitch, 2008). These hybrids may arise more readily via backcrossing or unidirectional hybridization, in which chromosome pairing is stabilized through resemblance to one parent, while chromosomes from the other parent may be partially lost, silenced, or excluded. Second, treating uniparentally homoploid hybrids as distinct from those that merely fall within the parental range but match neither parent (i.e., true intermediates) allows the index to preserve biologically meaningful patterns. These cases often have different evolutionary implications, particularly when hybrid swarms or introgression zones involve strong cytogenetic asymmetry. Without a directional correction, such biologically distinct configurations would be numerically indistinguishable from other midrange PDI values, thereby obscuring patterns of asymmetric hybridization and cytogenetic bias in large-scale datasets.

This treatment also facilitates clear classification of hybrid types in subsequent analyses, enabling researchers to test whether uniparental cytogenetic stabilization is disproportionately represented in artificial (horticultural) systems or in specific clades—insights that would be lost without directional encoding.

### Transformation

Chromosome number differences can span orders of magnitude across hybrid systems, especially when polyploidy is involved. To minimize the influence of these outliers and allow for meaningful comparison across a broad scale, we apply a logarithmic transformation to the absolute value of the signed index, restoring the original sign after transformation:

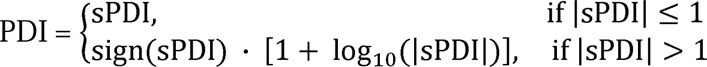

This transformation applied to the final index (PDI) compresses extreme values while preserving biologically meaningful distinctions at finer scales of chromosomal deviation.

### Compression and score interpretation

The above can be consolidated into a single formula represented below for ease of use:

*C_H_* = Hybrid chromosome number

*C_P1_*, *C_P2_* = Chromosome numbers of both parent species

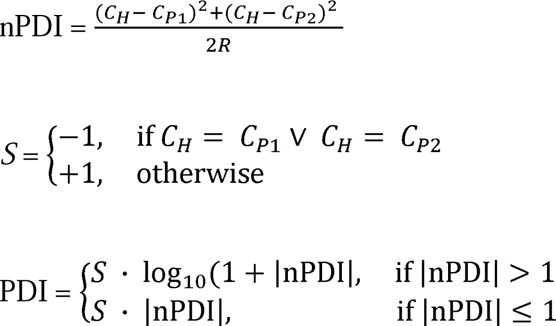

The final PDI scores are interpreted categorically to aid biological inference:

> (Category one): PDI = 0: Homoploid hybrid, where the hybrid matches both parental chromosome numbers.
>
> (Category two): –1 < PDI < 0: Uniparental homoploidy, where the hybrid matches one parent but not the other.
>
> (Category three): 0 < PDI ≤ 1: Intermediate, where the hybrid chromosome number is different from both parents but falls within their numerical range.
>
> (Category four): PDI > 1: Cytogenetically divergent or polyploid hybrid, where the hybrid chromosome number lies outside the parental range.

This framework allows the quantification of how “extreme” a hybrid’s chromosomal configuration is relative to its parents and enables downstream analysis of whether particular PDI classes are enriched under particular evolutionary conditions.

The PDI enables the analysis of large-scale patterns in cytogenetic compatibility across hybrid systems, providing a quantitative framework for detecting trends in the prevalence of homoploid, uniparentally homoploid, intermediate, and polyploid outcomes. By applying this metric to natural and horticultural hybrids, researchers can test whether certain chromosomal configurations are disproportionately represented, and whether cytogenetic divergence correlates with taxonomic group, life history, or hybrid origin. This approach facilitates new insights into the mechanisms underlying hybrid viability and stabilization, and opens the door to comparative analyses that assess the evolutionary repeatability of hybrid genome architecture. As such, the PDI offers a powerful tool for uncovering macroevolutionary patterns in plant hybridization, bridging cytogenetics, systematics, and evolutionary biology.

## Sampling

The Chromosome Counts Database (CCDB) (Rice et al., 2014) was used to find published chromosome counts for hybrid systems in all monocot families as defined by the Angiosperm Phylogeny Group (APG) (2016). For species with multiple citations on CCDB, the most recent of the original publications was chosen. Key words used to find hybrid papers outside of CCDB included the following: “[genus/family] ploidy,” “ [genus/family] chromosome counts,” “ [genus/family] chromosome number,” “karyotype,” “new [genus/family] hybrid species,” “hybridization in [genus/family],” and “[genus/family] cytology”. For publications with cytogenetic information solely for the hybrid species, up-to-date taxonomy and established parentage of each hybrid was confirmed using Kew’s Plants of the World Online (POWO, 2023). Chromosome counts for identified parents were determined by searching an appropriate subset of the previously listed key words.

Agricultural hybrids were excluded from analysis due to their long history—often exceeding a century—of intentional backcrossing (Vogel, 2009). In these breeding systems, artificial hybridization is typically used to achieve heterosis or hybrid vigor (Farinati et al., 2023), and repeated backcrossing is commonly employed to combine specific qualitative traits, such as enhanced disease resistance, while retaining other desirable characteristics (Poltronieri et al., 2020). Because many modern crop species have arisen primarily through such backcrossing (Farinati et al., 2023), their cytogenetic profiles are heavily shaped by human selection and would introduce bias into the analysis. In contrast, horticultural hybrids were included as backcrossing is generally less frequent and less extensive during the breeding of ornamental plants (Nomura et al., 2022). Hybrid systems lacking ploidy data for one or both parent species were excluded to ensure analytical consistency: The PDI calculation requires cytogenetic data from both parents, and attempting to calculate PDI scores without complete parental information would compromise the reliability of the meta-analysis. All data used in this study are available in Appendix S1.

## Statistical analysis

### Natural vs artificial hybrids

To visualize the distribution of PDI (Ploidy Deviation Index) scores among natural and artificial hybrids, we generated violin plots with embedded box-and-whisker plots using R (v4.3.0) with the ggplot2 package (Wickham, 2016). Natural hybrids are defined as taxa that arise without direct human intervention, whereas artificial hybrids are those produced through horticultural breeding programs. The violin plot approach was chosen to simultaneously display the kernel density of PDI scores and their median and interquartile ranges (Tukey, 1977), offering a robust visual comparison of central tendency and distributional spread.

To statistically evaluate whether the distributions of PDI scores differed significantly between natural and artificial hybrids, we performed an Anderson–Darling k-sample test, a nonparametric, distribution-free procedure that compares the entire distributions of two or more samples (Scholz and Stephens, 1987). This test is particularly appropriate here because it is sensitive to differences in the tails as well as the center of the distribution, and does not assume a specific parametric form of the underlying data — an assumption met by the heterogeneous and potentially non-normal distributions of PDI scores in hybrid populations.

### Comparisons in habit

Hybrid PDI scores were further compared among hybrids exhibiting five distinct growth habits: aquatic, epiphytic, evergreen, geophytic, and graminoid. Aquatic hybrids were defined as growing partially or fully submerged in water (Cook, 1990). Epiphytic hybrids were defined as growing on the surface of another plant without deriving nutrients from it (Benzing, 1990). Evergreen hybrids were operationally defined here as species with persistent leaves that do not fall into any of the other four habit categories; this is justified given that traditional secondary growth in monocots is rare or absent (Tomlinson and Zimmermann, 1969). Geophytes were defined as plants with periodic dieback of their above-ground structures while maintaining dormant underground stems (Dahlgren and Clifford, 1982), and graminoids were classified by their grass-like habit with typically narrow, parallel-veined leaves (Clayton and Renvoize, 1986). In cases where hybrids could reasonably be assigned to more than one habit category (for example, species exhibiting facultative aquatic and terrestrial growth), the primary habit was determined based on the predominant growth form reported in the literature or horticultural records, to ensure consistent and mutually exclusive classification.

The distribution of PDI scores for each habit was visualized using violin plots with embedded box-and-whisker plots in R (v4.3.0), providing both a summary of central tendency and the kernel density distribution across groups. Anderson–Darling k-sample tests were performed to evaluate significant differences among the five habit types, chosen for their sensitivity to differences in the entire distribution, including the tails, and because they do not assume a parametric distribution of the data (Scholz and Stephens, 1987). Because the overall test indicated significant differences among the five habits, pairwise Anderson–Darling tests were then conducted to identify which specific pairs of habits differed significantly.

For each pairwise comparison of habits found to be significant, a multinomial logistic regression was used to evaluate whether the predicted probabilities of PDI categories varied by habit type. For this analysis, PDI scores were binned into four biologically meaningful categories: less than zero (uniparentally homoploid), equal to zero (homoploid), between zero and one (intermediate), and greater than one (polyploid). The multinomial regression was performed using the nnet package (Ripley, 2022), with estimated marginal means (emmeans) contrasts subsequently tested to assess the significance of pairwise predicted probabilities across categories (Lenth, 2022). Finally, for each significant pairwise difference, the means of PDI scores within each hybrid category were visualized with bar plots including standard error bars, and tested for statistical significance using a Mann–Whitney U test. These nonparametric tests are appropriate because the data did not meet assumptions of normality and included ordinal or non-continuous categories, allowing robust inferences without parametric assumptions about their distributions.

### Comparisons in hybrid genus size and genus chromosomal range

To investigate the relationship between hybrid cytogenetic patterns and broader genus-level characteristics, we compared values of “genus size” and “genus chromosome range” across PDI categories. Genus size was quantified as the number of accepted species per genus using data from Plants of the World Online (Kew Science, 2023). Genus chromosome range was calculated by subtracting the smallest total chromosome count of any species in the genus from the largest, using records compiled from the Chromosome Counts Database (CCDB; Rice et al., 2014). To ensure consistency, intergeneric hybrids were excluded from these analyses.

As in previous sections, PDI scores were binned into four biologically meaningful categories (uniparentally homoploid, homoploid, intermediate, and polyploid) and compared across groups. The distribution of genus size and genus chromosome range within each hybrid condition was visualized using violin plots with embedded box-and-whisker plots, summarizing both central tendency and kernel density. Anderson–Darling k-sample tests were performed to test whether distributions of genus size or chromosome range differed significantly across hybrid conditions.

To further test for significant differences between hybrid conditions, mean values with standard error bars were visualized using bar plots, and the Mann–Whitney U test was applied to assess whether average genus size or genus chromosome range differed between categories. Finally, to explore relationships between genus size or genus chromosome range and PDI scores within each hybrid condition, scatter plots were generated with ordinary least squares regression lines added for visualization. Although these regression lines illustrate potential trends, the formal assessment of association was carried out using Kendall’s tau correlation, which is well-suited to non-normal data and tests for monotonic associations without assuming linearity (Gibbons and Chakraborti, 2020).

## RESULTS

### The use of the ploidy deviation index

The ploidy deviation index (PDI) effectively classified each hybrid system into consistent, biologically meaningful categories across the dataset. Hybrids with chromosome numbers identical to both parents were assigned to the homoploid category (PDI = 0). Those with a chromosome number identical to only one parent were categorized as uniparentally homoploid, showing negative PDI values. Hybrids with chromosome numbers intermediate between both parents (0 < PDI ≤ 1) were classified as intermediate, while those exceeding the parental chromosome range (PDI > 1) were categorized as polyploid. This scheme was consistently applicable to both natural and horticultural hybrids, providing a standardized approach to comparing chromosome numbers among diverse hybrid systems.

Within these categories, homoploid hybrids consistently scored exactly zero, indicating complete chromosome-number matches to both parents. Uniparentally homoploid hybrids displayed negative PDI values ranging from –0.1 to –1.0, with more negative scores reflecting a greater proportional deviation from the non-matching parent’s chromosome number. Intermediate hybrids showed PDI values between 0.1 and 1.0, with higher intermediate scores indicating a hybrid chromosome number more equally distant from both parents, while lower intermediate scores closer to 0.1 reflected hybrids whose chromosome numbers were only slightly shifted from one parent but still fell within the parental range. Polyploid hybrids had positive PDI scores exceeding 1, ranging up to 4.5, with higher values reflecting progressively larger deviations from the parental cytotypes and increased multiples of the parental chromosome complements. These distributions were clearly separated across categories, with no observed overlap in score assignments (see Figure S1).

### Natural vs artificial hybrids

The distribution of PDI scores differed between natural and artificial hybrids (Figure 1). For natural hybrids (n = 135), PDI scores ranged from –1.0 to 4.07, with a mean of 0.49, median of 0.0, and an interquartile range (IQR) of 0.5. For artificial hybrids (n = 56), PDI scores ranged from –1.0 to 4.36, with a mean of 0.67, median of 0.375, and an interquartile range (IQR) of 0.5. An Anderson–Darling k-sample test was conducted to evaluate whether the overall distributions differed significantly between natural and artificial hybrids. The test statistic was T.AD = 0.60675 with a p-value of 0.18589, indicating no statistically significant difference in the distribution of PDI scores between the two groups (Figure 1).

**Figure 1.**
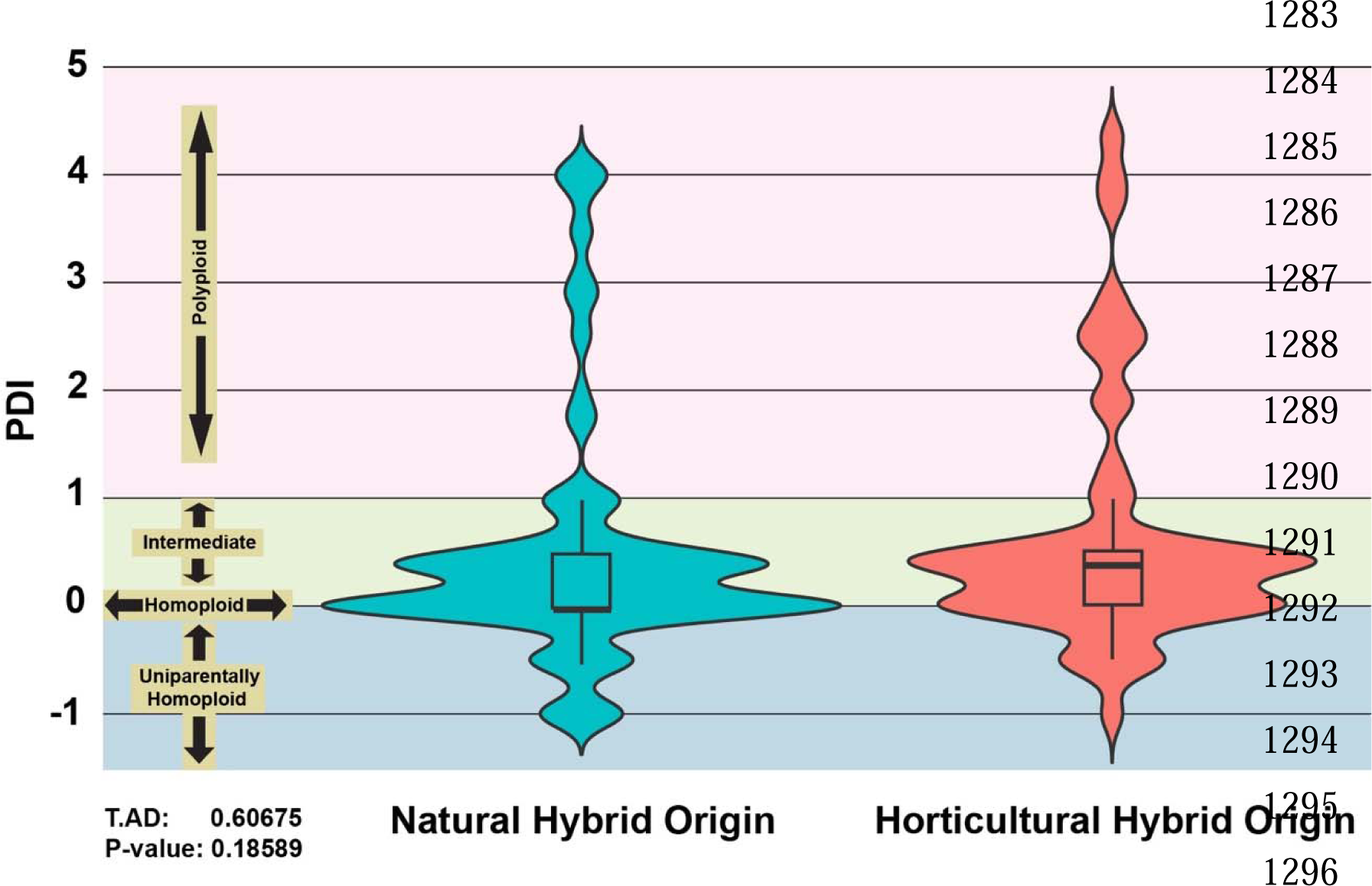
Violin plots with box and whisker overlay illustrating the distribution of PDI scores of hybrids resulting from natural (left; n = 135) vs horticultural (right; n = 56) processes. PDI score range corresponding to hybrid condition (uniparentally homoploid, homoploid, intermediate, polyploid) outlined on the left. Anderson-Darling k-sample test summary outlined in the lower left; T.AD = Anderson-Darling test statistic.

### Comparisons in habit

The distribution of PDI scores was summarized across five habit types (Figure 2). Among aquatic hybrids (n = 36), PDI scores ranged from –1.0 to 4.01, with a mean of 0.13, a median of 0.0, and an interquartile range (IQR) of 0.375. Epiphytic hybrids (n = 11) showed PDI scores from –1.0 to 3.73, with a mean of 0.96, a median of 0.5, and an IQR of 1.28. Evergreen hybrids (n = 8) had PDI scores between –1.0 and 0.0, with a mean of –0.19, median of 0.0, and IQR of 0.125. Geophytic hybrids (n = 88) ranged from –1.0 to 4.36, with a mean of 0.65, median of 0.375, and IQR of 0.625. Graminoid hybrids (n = 36) showed PDI scores from –1.0 to 4.07, with a mean of 0.50, median of 0.375, and IQR of 0.52.

**Figure 2.**
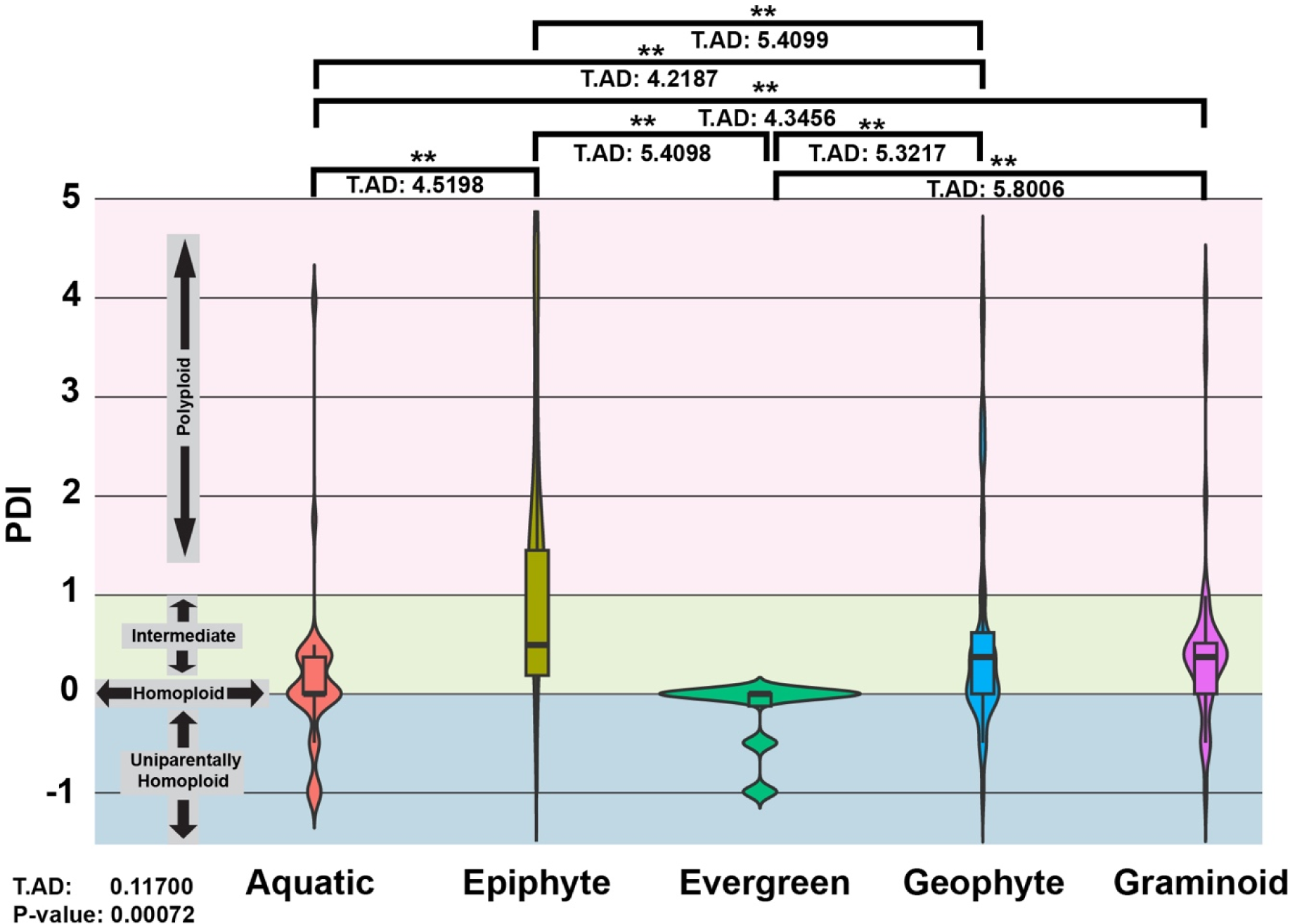
Violin plots with box and whisker overlay illustrating the distribution of PDI scores of hybrids resulting from various habits: aquatic (n = 36), epiphyte (n = 11), evergreen (n = 9), geophyte (n = 88), and graminoid (n = 36). PDI score range corresponding to hybrid condition (uniparentally homoploid, homoploid, intermediate, polyploid) outlined on the left. Anderson-Darling k-sample test summary comparing the overall distribution of all five habits outlined in the lower left; T.AD = Anderson-Darling test statistic. Significant (P < 0.05) pairwise comparisons between habits are shown by brackets with P-value above represented by an asterisk above (* = P < 0.05; ** = P < 0.005) and Anderson-Darling test statistic underneath.

An Anderson–Darling k-sample test indicated significant differences among the overall distributions of PDI scores by habit (Figure 2: T.AD = 0.11700, p =0.00072). Pairwise Anderson–Darling tests revealed seven significant comparisons in PDI score distribution: aquatic vs. epiphyte (T.AD = 4.5198, p < 0.005), aquatic vs. geophyte (T.AD = 4.2187, p < 0.005), aquatic vs. graminoid (T.AD = 4.3456, p < 0.005), epiphyte vs. evergreen (T.AD = 5.4098, p < 0.005), epiphyte vs. geophyte (T.AD = 5.4099, p < 0.005), evergreen vs. geophyte (T.AD = 5.3217, p < 0.005), and evergreen vs. graminoid (T.AD = 5.8006, p < 0.005) (Figure 2).

A multinomial logistic regression was conducted to examine the predicted probabilities of hybrid conditions across the seven significant habit pairs identified by the Anderson–Darling tests (Figure 3). Eight pairwise comparisons showed significant differences in predicted probabilities. Geophytic hybrids had a significantly greater predicted probability of being classified as polyploid compared to aquatic hybrids (estimated difference in predicted probability (EST): – 0.138; standard error (SE): 0.057; p < 0.05). Aquatic hybrids had a significantly greater predicted probability of being homoploid compared to graminoid hybrids (EST: 0.306, SE: 0.101, p < 0.05), while graminoids were significantly more likely to be intermediate than aquatics (EST: – 0.333, SE: 0.111, p < 0.05).

**Figure 3.**
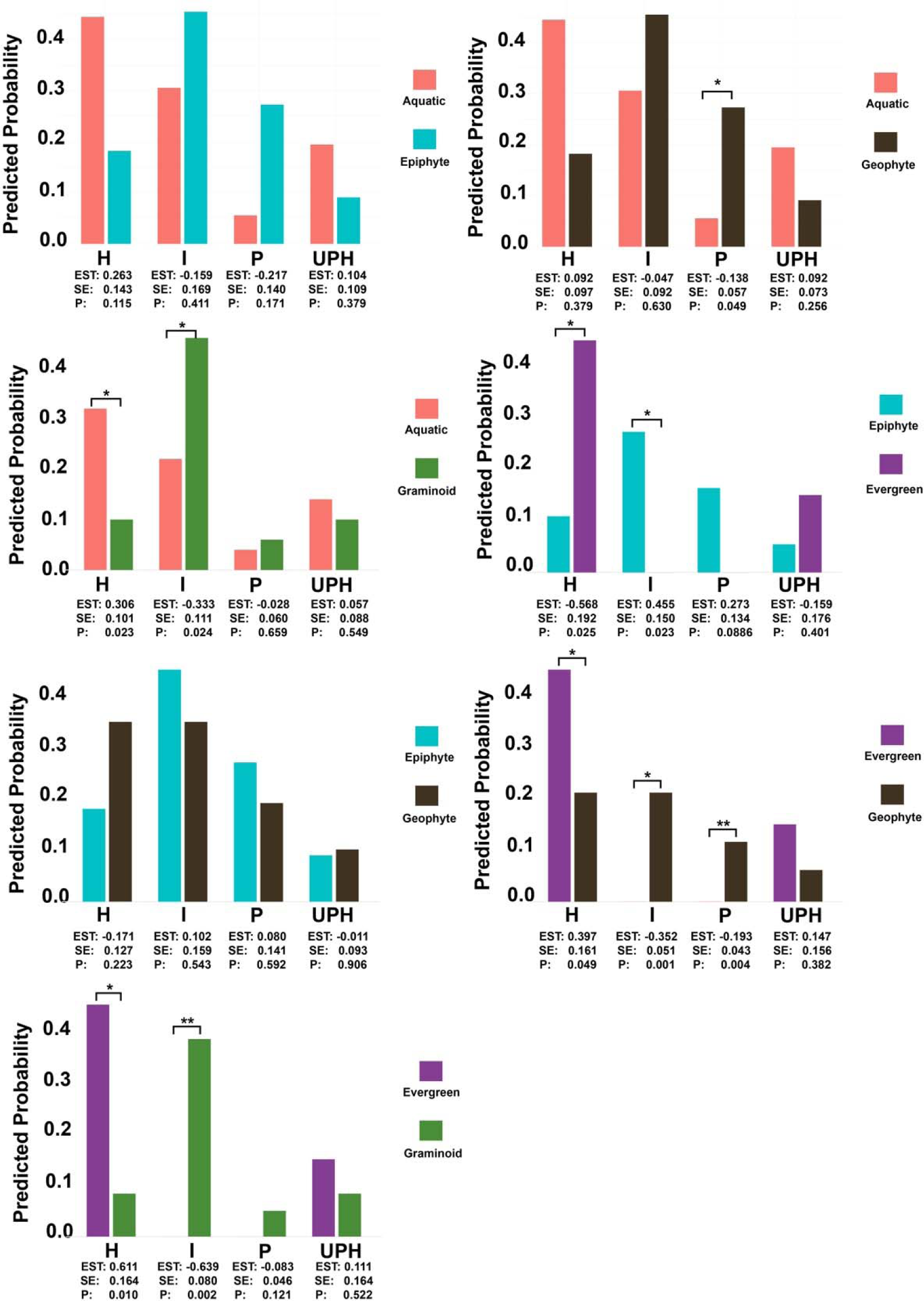
Multinomial regression with estimated marginal means illustrating differences in predicted probability of each hybrid condition (homoploid (H), intermediate (I), polyploid (P), uniparentally homoploid (UPH)) for pairwise comparisons of habit that exhibit significant differences in PDI score distribution. Significant (P < 0.05) pairwise comparisons between habits are shown by brackets with P-value above represented by an asterisk above (* = P < 0.05; ** = P < 0.005). EST = estimated difference in predicted probability; SE = standard error; P = P-value.

Epiphytic, geophytic, and graminoid hybrids were found to be significantly more likely to be intermediate than evergreen hybrids (EST: 0.455, –0.352, –0.639; p < 0.05, p < 0.5, p < 0.005, respectively); however, it should be noted that no intermediate hybrids were recorded for the evergreen category (Figure 3; purple). In contrast, evergreen hybrids were significantly more likely to be classified as homoploid than epiphytic, geophytic, and graminoid hybrids (EST: – 0.568, 0.397, 0.611, respectively; all p < 0.5), though the sample size for evergreen hybrids was limited.

Mean PDI scores within each hybrid condition were again compared for the seven significant pairwise habit comparisons identified above (Figure 4). Among these, only one comparison showed a statistically significant difference: Geophytic intermediate hybrids had significantly higher PDI scores than aquatic intermediate hybrids based on the Mann–Whitney U test (W = 130.5, p < 0.05) (Figure 4; pink v. black). No other pairwise comparisons of mean PDI scores within hybrid conditions were significant.

**Figure 4.**
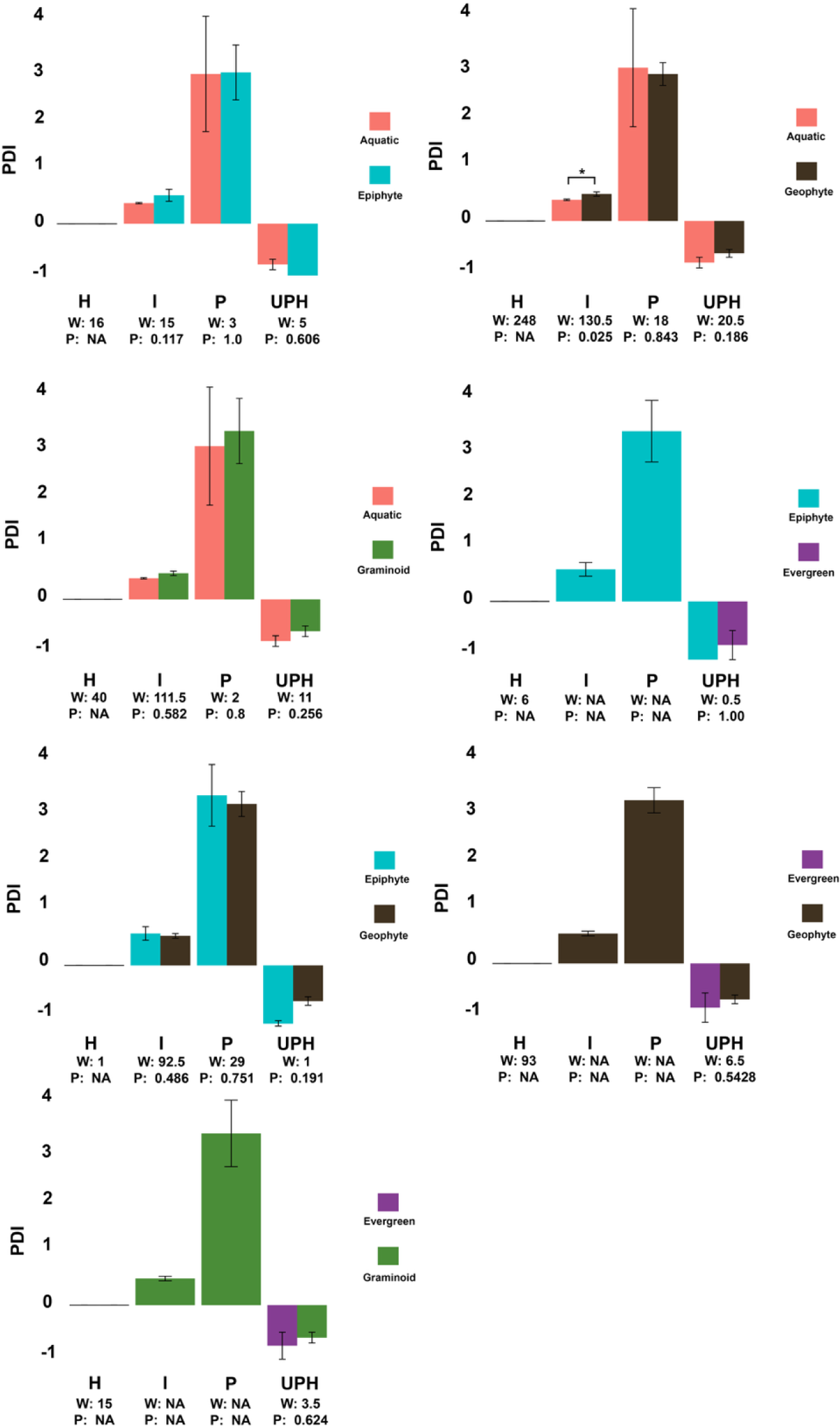
Comparison of mean PDI scores (with standard error) within each hybrid condition (homoploid (H), intermediate (I), polyploid (P), uniparentally homoploid (UPH)) for pairwise comparisons of habit that exhibit significant differences in PDI score distribution. Significant (P < 0.05) pairwise comparisons between habits, determined by the Mann-Whitney U-test, are shown by brackets with P-value above represented by an asterisk above (* = P < 0.05; ** = P < 0.005). W = Mann-Whitney test statistic; P = P-value. Bars are doubled in width when the sample size is zero for a given hybrid condition within a habit type. Mann-Whitney test summary reads “NA” if the sample size was insufficient or if all PDI values are equivalent (such as for homoploid hybrids).

### Comparisons in hybrid genus size

The distribution of genus sizes was summarized across the four hybrid conditions (Figure 5). Homoploid hybrids (n = 65) had genus sizes ranging from 7 to 1089 species, with a mean of 95.1, median of 77, and an interquartile range (IQR) of 56. Intermediate hybrids (n = 70) showed sizes of genera ranging from 13 to 2094 species, with a mean of 602.4, median of 90, and an IQR of 519.5. Polyploid hybrids (n = 29) were found in genera that ranged from 1 to 1600 species, with a mean of 174.3, median of 76, and an IQR of 14. Uniparentally homoploid hybrids (n = 24) were in genera with sizes from 24 to 2094 species, with a mean of 300.5, median of 83, and an IQR of 38 (Figure 5A).

**Figure 5.**
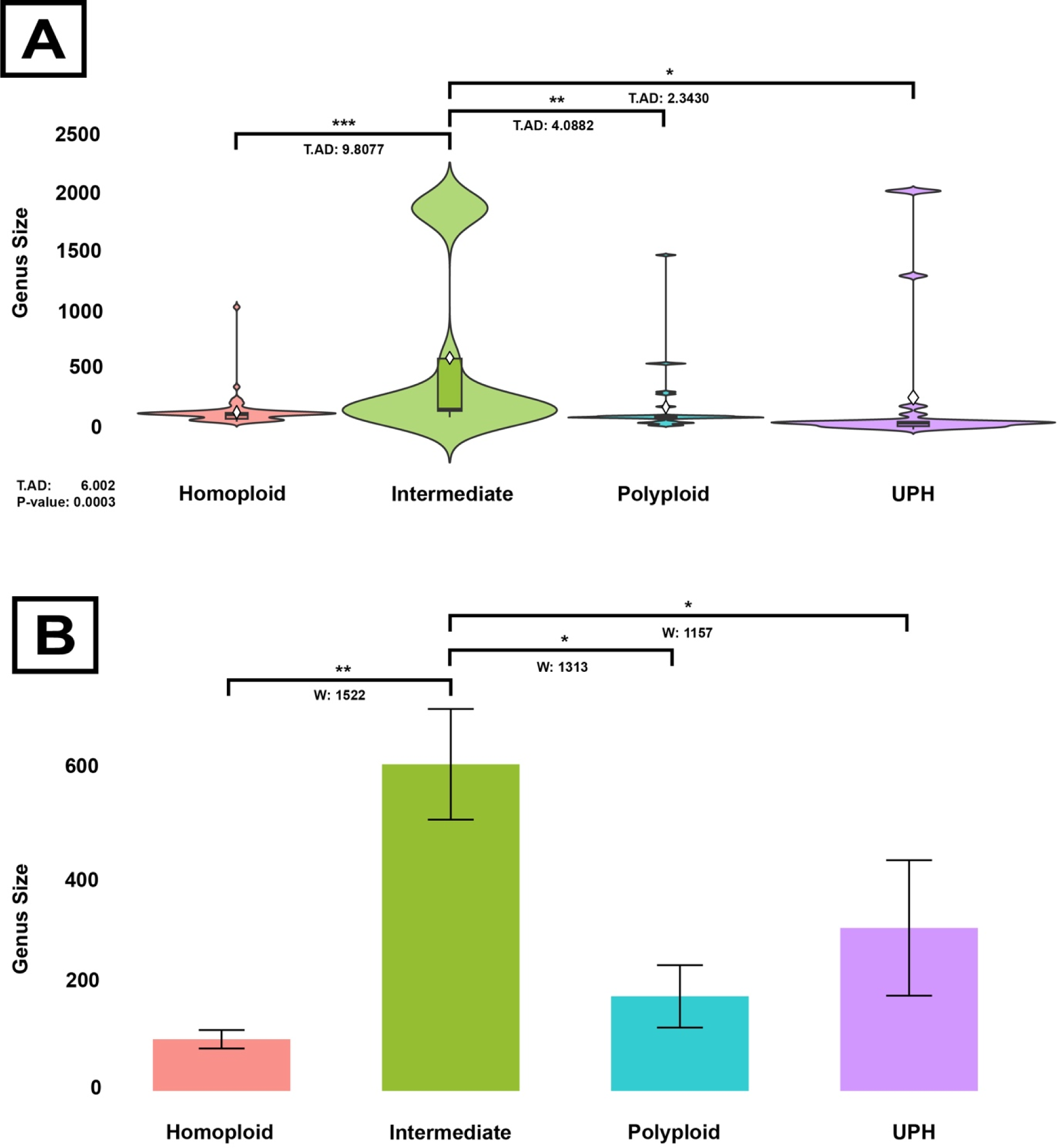
Comparisons of genus size and occurrence of each hybrid condition (homoploid (n = 65), intermediate (n = 70), polyploid (n = 32), uniparentally homoploid (UPH) (n = 24)). **A.** Violin plots with box and whisker overlay illustrating the distribution of genus sizes within each hybrid condition. Anderson-Darling k-sample test summary comparing the overall distribution of all four conditions outlined in the lower left; T.AD = Anderson-Darling test statistic. Significant (P < 0.05) pairwise comparisons between habits are shown by brackets with P-value above represented by an asterisk above (* = P < 0.05; ** = P < 0.005, *** = P < 0.0005) and Anderson-Darling test statistic underneath. **B.** Comparison of mean genus size (with standard error) within each hybrid condition. Significant (P < 0.05) pairwise comparisons between conditions, determined by the Mann-Whitney U-test, are shown by brackets with P-value above represented by an asterisk above (* = P < 0.05; ** = P < 0.005) and Mann-Whitney test statistic (W) below.

An Anderson–Darling k-sample test revealed a significant difference in the distribution of genus sizes among the four hybrid conditions (Figure 5B; T.AD = 6.002, p < 0.0005). Three significant pairwise comparisons were identified: intermediate vs. homoploid (T.AD = 9.8077, p < 0.0005), intermediate vs. polyploid (T.AD = 4.0882, p < 0.005), and intermediate vs. uniparentally homoploid (T.AD = 2.430, p < 0.05).

In addition, mean sizes of the genera were compared between hybrid conditions using Mann–Whitney U tests. Three significant pairwise comparisons emerged: intermediate hybrids had significantly larger mean genus sizes than homoploid (W = 1522, p > 0.005), polyploid (W = 1313, p < 0.05), and UPH hybrids (W = 1157, p < 0.05) (Figure 5).

Scatter plots with regression lines were generated to visualize the relationship between PDI scores and genus size for each of the four hybrid conditions (Figure 6). Kendall’s tau rank correlation tests were applied to formally assess the association in each category. Among these, only polyploid hybrids showed a significant correlation between PDI score and genus size, with larger genera associated with higher PDI values (Figure 6C; Kendall’s tau test statistic (τ) = 0.332, p < 0.05). No significant correlations were detected between PDI score and genus size in the homoploid, intermediate, or uniparentally homoploid hybrid categories (Figure 6A,B,D).

**Figure 6.**
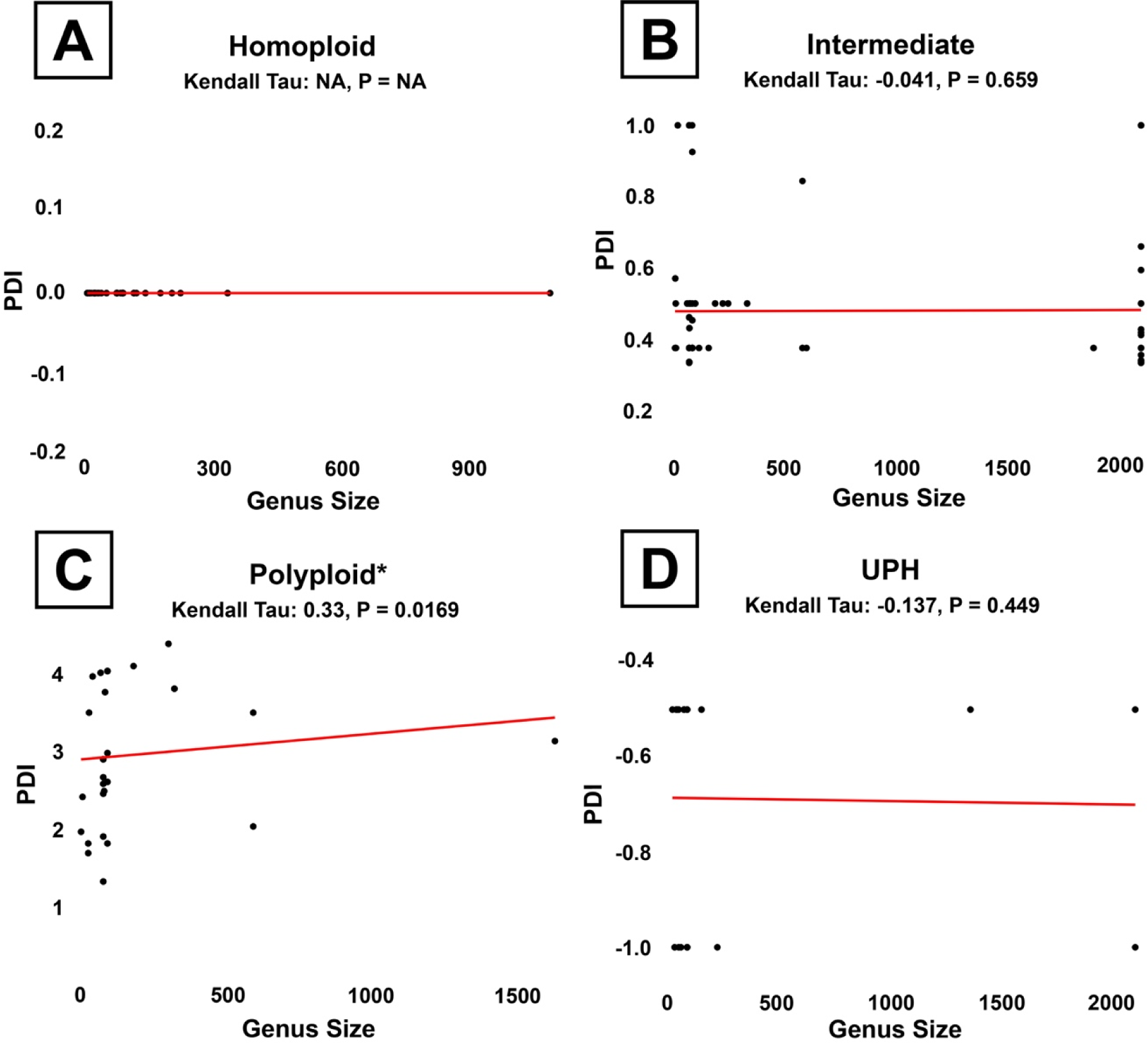
Scatter plots with ordinary least squares regression line comparing the correlation of PDI score and genus size within each hybrid condition **A.** Homoploid (n = 65). **B.** Intermediate (n = 70). **C.** Polyploid (n = 32) **D.** Uniparentally homoploid (UPH) (n = 24). Summary statistics from the Kendall’s tau rank correlation tests are above each plot; Kendall Tau = Kendall’s tau rank correlation test statistic; P = P-value. Significant (P < 0.05) correlations are marked by an asterisk.

### Comparisons in genus chromosomal range

The distribution of hybrid genus chromosome ranges was summarized across the four hybrid conditions (Figure 7). Homoploid hybrids (n = 65) had ranges of chromosome number across their genus from 0 to 110, with a mean of 34.1, median of 31, and an interquartile range (IQR) of 22. Intermediate hybrids (n = 70) were found to be in genera that exhibited chromosome number ranges between 21 and 242, with a mean of 89.1, median of 45, and an IQR of 96. Polyploid hybrids (n = 28) had genus ranges from 13 to 119, with a mean of 42.3, median of 39.5, and an IQR of 22.5. Uniparentally homoploid hybrids n = 24) showed genus ranges from 15 to 242, with a mean of 62.7, median of 45, and an IQR of 36.8.

**Figure 7.**
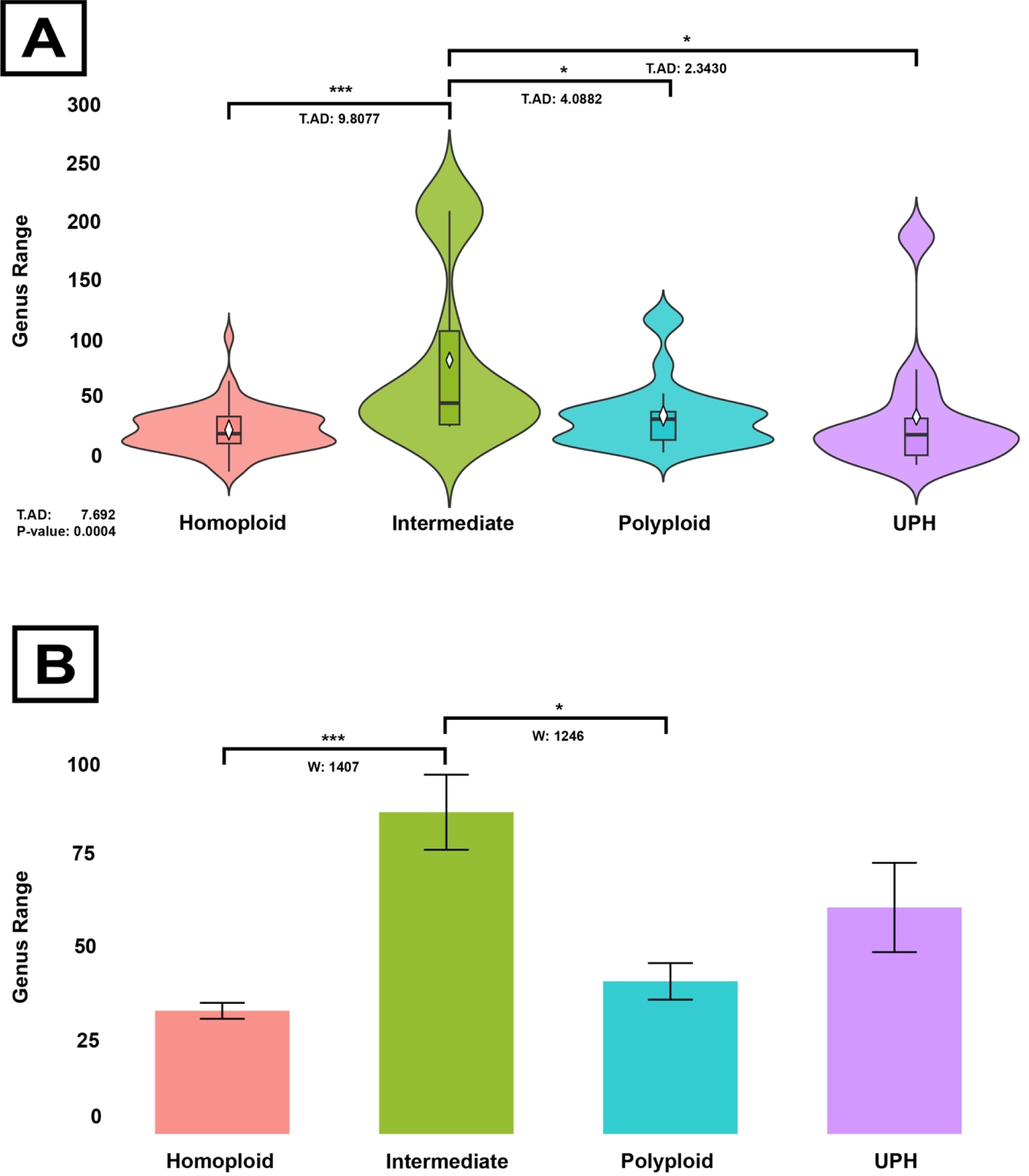
Comparisons of genus chromosomal range and occurrence of each hybrid condition (homoploid (n = 65), intermediate (n = 70), polyploid (n = 32), uniparentally homoploid (UPH) (n = 24)). **A.** Violin plots with box and whisker overlay illustrating the distribution of genus chromosomal range within each hybrid condition. Anderson-Darling k-sample test summary comparing the overall distribution of all four conditions outlined in the lower left; T.AD = Anderson-Darling test statistic. Significant (P < 0.05) pairwise comparisons between habits are shown by brackets with P-value above represented by an asterisk above (* = P < 0.05; ** = P < 0.005, *** = P < 0.0005) and Anderson-Darling test statistic underneath. **B.** Comparison of mean genus chromosomal range (with standard error) within each hybrid condition. Significant (P < 0.05) pairwise comparisons between conditions, determined by the Mann-Whitney U-test, are shown by brackets with P-value above represented by an asterisk above (* = P < 0.05; ** = P < 0.005) and Mann-Whitney test statistic (W) below.

An Anderson–Darling k-sample test indicated significant differences in the distribution of genus chromosome ranges among the four hybrid categories (Figure 7A; T.AD = 7.692, p < 0.0005). Three significant pairwise comparisons were detected: intermediate vs. homoploid (T.AD = 9.8077, p < 0.0005), intermediate vs. polyploid (T.AD = 4.0882, p < 0.05), and intermediate vs. uniparentally homoploid (T.AD = 2.3430, p < 0.05).

Mean genus chromosome ranges were also compared between hybrid conditions using Mann–Whitney U tests (Figure 7B). Intermediate hybrids were found to have significantly higher mean genus chromosome ranges than homoploid (W = 1407, p < 0.0005) and polyploid hybrids (W = 1246, p < 0.05).

Scatter plots with regression lines were generated to visualize the relationship between PDI scores and genus chromosome range within each of the four hybrid conditions (Figure 8). Kendall’s tau rank correlation tests were performed to formally assess these associations. Among the four conditions, only polyploid hybrids showed a statistically significant correlation between PDI score and genus chromosome range (Figure 8C; Kendall’s tau test statistic (τ) = 0.341, p < 0.05), indicating that higher PDI values tended to be associated with greater genus chromosome range. Like above, no significant correlations were detected in homoploid, intermediate, or uniparentally homoploid hybrids (Figure 8A,B,D).

**Figure 8.**
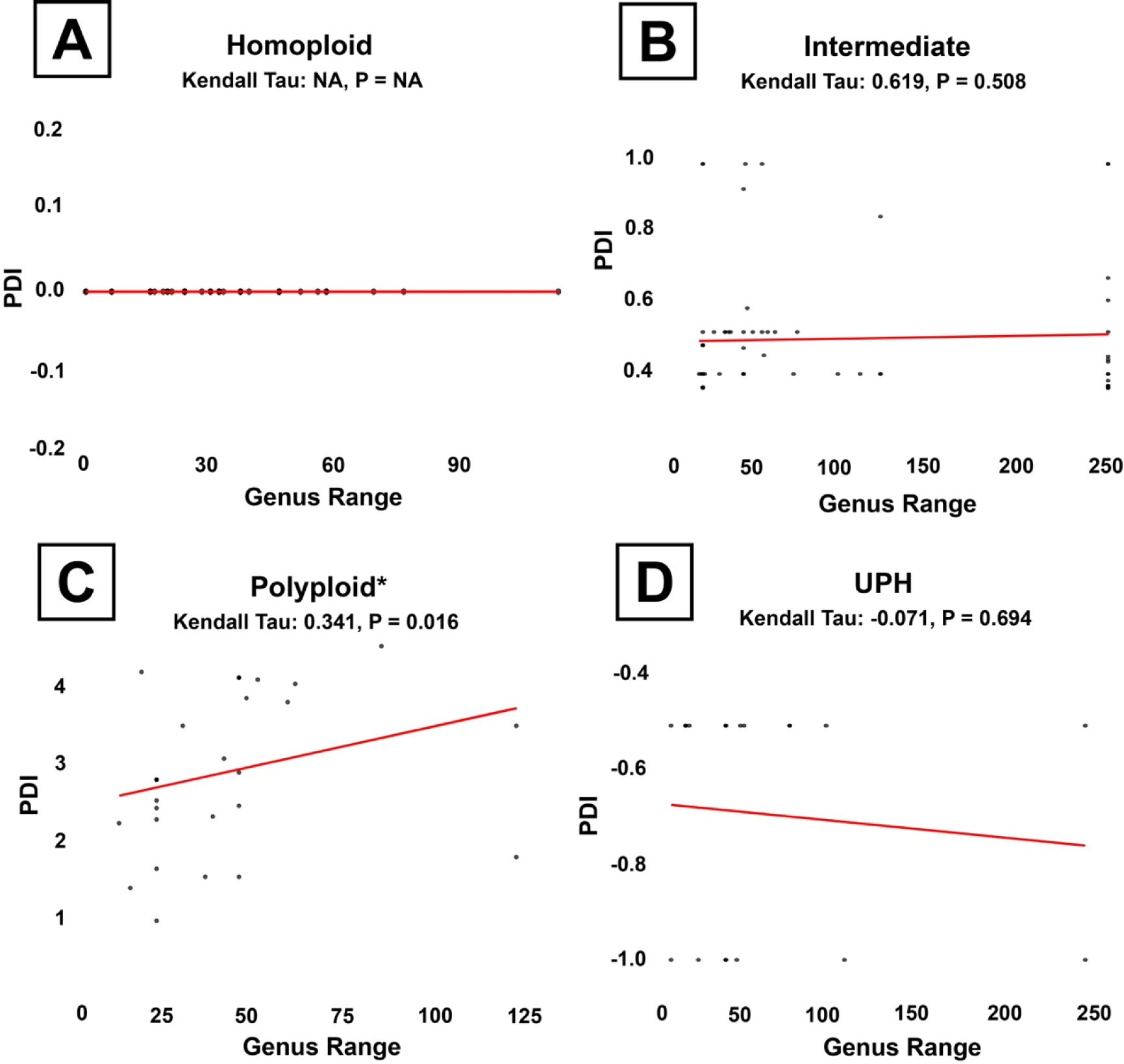
Scatter plots with ordinary least squares regression line comparing the correlation of PDI score and genus chromosomal range within each hybrid condition **A.** Homoploid (n = 65). **B.** Intermediate (n = 70). **C.** Polyploid (n = 32) **D.** Uniparentally homoploid (UPH) (n = 24). Summary statistics from the Kendall’s tau rank correlation tests are above each plot; Kendall Tau = Kendall’s tau rank correlation test statistic; P = P-value. Significant (P < 0.05) correlations are marked by an asterisk.

## DISCUSSION

### Ploidy deviation index performance

In this study, the ploidy deviation index (PDI) enables a more biologically meaningful analysis of chromosomal divergence in monocot hybrids than traditional binary classifications allow. By quantifying both the magnitude and direction of chromosome-number differences, the PDI revealed the potential influence of numerous biological variables, patterns of cytogenetic asymmetry, and tolerance of chromosomal shifts across hundreds of hybrid systems. Importantly, because the PDI can be interpreted in well-defined categories (homoploid, uniparentally homoploid, intermediate, and polyploid), it supports both continuous and categorical analyses, allowing flexible but rigorous testing of hybrid cytogenetic patterns. This framework captured variation in hybrid chromosome architecture that would have been obscured by simpler classifications, permitting the dataset to be interpreted with greater evolutionary and biological resolution.

### The frequency of homoploid hybrids in natural and artificial systems

Using a meta-analysis approach, this study examines how hybrid cytogenetic divergence varies across different hybrid contexts. One of the most striking findings was the overall high frequency of homoploid hybrids in the dataset, far exceeding what might be expected from the literature, which has historically emphasized the rarity of homoploid hybrid speciation (Rieseberg, 1997; Gross and Rieseberg, 2005; Abbott et al., 2013). While most hybrids included here have not been tested for reproductive isolation or formal species status, their chromosome-level compatibility suggests that homoploid hybrid formation is far from uncommon. These results indicate no clear pattern of greater polyploid hybrid formation compared to homoploid hybrids across monocots.

In fact, many of the analyses presented here — including comparisons across hybrid contexts and growth habits — suggest the opposite, challenging long-standing claims that polyploid hybrid species presumably dramatically outnumber homoploid ones (Stebbins, 1950; Soltis and Soltis, 1999; Wood et al., 2009). This raises doubts as to whether the high perceived frequency of polyploid hybrid species truly reflects a biological reality, or instead results from a bias in recognizing homoploid hybrids due to the lack of obvious postzygotic barriers or species status confirmation (Mallet, 2007; Feliner et al., 2017).

The comparison between natural and artificial hybrids offers an opportunity to further test this question. It has often been argued that homoploid hybrids rarely persist in natural systems because they lack sufficient reproductive isolation from their parental species, making them vulnerable to repeated backcrossing or genetic swamping (Grant, 1981; Rieseberg, 1997). By contrast, artificial systems — particularly horticultural breeding programs — can maintain homoploid hybrids through human-controlled reproductive barriers, effectively preserving them in cultivation even if they would fail to persist in the wild. Based on this reasoning, we expected a higher proportion of homoploid hybrids in artificial systems relative to natural ones, reflecting the presumed inability of homoploid hybrids to establish as independent lineages under natural evolutionary processes.

However, the results of this study did not support that prediction. There was no significant difference in the distribution of hybrid conditions between natural and artificial systems, suggesting that the relative proportions of homoploid, intermediate, and polyploid hybrids are roughly equivalent across both contexts (see Figure 1). At least among monocot taxa, these findings imply that homoploid hybrids may be more evolutionarily stable and common than previously assumed, challenging the notion that reproductive isolation is an insurmountable barrier to their persistence. Instead, the comparable frequency of homoploid hybrids in both natural and artificial systems raises the possibility that ecological factors or ongoing selection may help maintain these hybrids in nature, even in the absence of strong postzygotic reproductive barriers.

### Differences in growth habit

Examining differences in PDI across growth habits offers valuable insight into whether ecological context might structure patterns of hybrid cytogenetic divergence. Distinct growth forms can reflect differences in life history, environmental stability, and opportunities for hybrid establishment. For example, aquatic or geophytic taxa may experience more frequent disturbance or habitat turnover (Cook, 1990; Benzing, 1990), potentially influencing the success of hybrids with varying chromosomal architectures, while epiphytes and evergreen taxa may be subject to more stable but spatially restricted conditions (Benzing, 1990; Tomlinson and Zimmermann, 1969). Investigating these patterns can therefore illuminate whether certain ecological strategies facilitate or constrain particular cytogenetic hybrid types.

In this study, PDI distributions differed significantly among growth habits, revealing that geophytes and epiphytes were more likely to harbor intermediate or polyploid hybrids, while aquatics showed a higher proportion of homoploid hybrids (see Figures 2,3). Evergreen hybrids appeared strongly biased toward homoploid or uniparentally homoploid outcomes; however, no intermediate hybrids were recorded for evergreen taxa, and their overall sample size was very small. Although the Anderson–Darling and multinomial tests adjust for small sample sizes, drawing firm conclusions about evergreen taxa will require additional data from hybrid systems exhibiting this growth habit.

Focusing on the comparisons with more robust sample sizes, the observed differences suggest that habitat features could shape hybridization outcomes. Aquatic hybrids, for example, were significantly more likely to be homoploid compared to graminoids and less likely to be polyploid than geophytes (Figure 3). This pattern is consistent with the idea that aquatic systems, which often feature stable water-dispersed propagules and clonal/vegetative reproduction, may buffer hybrid lineages from the need for dramatic chromosomal change to maintain persistence (Les.

1988; Barrett, 2015). In aquatic environments, vegetative propagation and fragmentation can allow hybrids to persist and spread even if sexual reproduction with parent species is genetically possible, reducing selective pressure for chromosome doubling or other cytogenetic restructuring (Philbrick and Les, 1996; Santamaría, 2002). Supporting this, intermediate hybrids in aquatic systems showed significantly lower PDI scores than intermediate hybrids in geophytic systems (Figure 3; pink v. black), suggesting that even when hybrid chromosomal shifts occur in aquatic habitats, they tend to be more conservative and closer to parental karyotypes, likely through backcrossing. As a result, homoploid and moderately divergent intermediate hybrids may more readily persist and establish in aquatic habitats compared to more terrestrial systems, where sexual reproduction with parental species may pose a greater obstacle to persistence.

### The potential role of genus size and chromosomal range

Beyond growth habit, patterns of hybrid cytogenetic divergence were also evaluated in relation to the size of the corresponding genus (genus size) and the range in number of chromosomes found across that genus (genus chromosome range). Monocot genera show striking differences in species richness, ranging from monotypic genera such as *Orontium* L. (Araceae) to hyper-diverse lineages like *Carex* L. (Cyperaceae) which exceeds 2,000 species (Global Carex Group, 2015). This diversity implies that larger genera likely encompass more divergent evolutionary lineages with broader chromosomal variation, while smaller genera may retain more uniform karyotypes and stronger reproductive boundaries (Levin, 2002; Weiss-Schneeweiss and Schneeweiss, 2013). As a result, genus size acts as a useful proxy for assessing the scope of potential cytogenetic barriers and opportunities in hybridization.

In this dataset, the intermediate hybrid category became progressively more frequent as the genus size increased. Intermediate hybrids ultimately represented the most common category of hybrid documented in each of the largest genera sampled. This suggests that intermediate chromosomal shifts may best accommodate the cytogenetic disparity present among more distantly related parental species in the more species-rich genera, where perfect homoploidy is less feasible but moderate shifts still allow sufficient meiotic pairing and stability (Ramsey and Schemske, 1998; Levin, 2002). Notably, however, this correlation raises the question of directionality—whether large genus size (i.e. having a higher number of species) facilitates more hybridization, or whether frequent hybridization has itself contributed to the expansion of genus size over evolutionary time. Several studies have proposed that hybridization can promote taxonomic diversification by generating novel lineages and facilitating ecological niche expansion (Rieseberg, 1997; Mallet, 2007; Soltis and Soltis, 2009), suggesting that hybridization may not merely be tolerated in large genera but could actively drive their diversification. In this sense, the prevalence of intermediate hybrids in larger genera may reflect both a greater opportunity for hybridization due to broader karyotypic and ecological diversity, and a possible legacy of past hybrid events that have contributed to the radiation and species richness of the genus. In these systems, when polyploid hybrids do occur, they often exhibit extremely high PDI values, reflecting substantial cytogenetic divergence from their parents. Polyploidization likely provides a robust mechanism for restoring meiotic stability in such cases by doubling the genome, allowing homologous pairing to proceed despite pronounced karyotypic differences (Soltis and Soltis, 1999; Otto and Whitton, 2000; Madlung, 2013). Thus, in large and chromosomally diverse genera, intermediate cytogenetic configurations may represent a common and evolutionarily productive outcome of hybridization, while polyploidy serves as a more radical yet effective solution to extreme chromosomal mismatches (Wood et al., 2009; Soltis et al., 2014).

To complement the genus size analysis, we also explicitly tested the effect of genus chromosome range, which is known to correlate strongly with genus size in monocots and other angiosperms (Rice et al., 2014). The results broadly mirrored those observed for genus size: intermediate hybrids were associated with genera exhibiting far greater chromosome number ranges on average than homoploid or polyploid hybrids, while uniparentally homoploid hybrids fell closer to the midrange. This further suggests that intermediate cytogenetic configurations are particularly favored in genera with broad chromosomal diversity, where hybrid formation must reconcile moderate to large parental differences in the absence of genome duplication.

As with genus size, PDI values were also found to increase within polyploid hybrids as genus chromosome range increased. This provides additional evidence that in genera characterized by high chromosomal variability, polyploid hybrids tend to arise with greater cytogenetic divergence from their parents. In these cases, polyploidy again appears to serve as a stabilizing mechanism to overcome extreme karyotypic mismatches by providing duplicate homologs for pairing during meiosis (Soltis and Soltis, 1999; Otto and Whitton, 2000; Madlung, 2013). Together, these patterns reinforce the conclusion that genera with greater chromosomal and taxonomic diversity promote intermediate chromosome numbers in hybrids and, when they occur, highly divergent polyploid configurations, likely depending on the scale of parental cytogenetic differences.

### Limitations

One limitation of this study is the restricted sample size of hybrid systems with fully documented parental chromosome counts. While the dataset assembled here represents one of the most comprehensive efforts to date, its scope is ultimately constrained by the limited availability of cytogenetic data across monocots. Much of the foundational karyotype literature was published between 1982 and 1986 (see Figure S2), with a sharp decline in new chromosome count reports in the decades since. This decrease in traditional karyotype work is likely due to shifting research priorities toward molecular markers and genomic sequencing approaches, as well as a loss of taxonomic expertise in classical cytogenetics. As a result, many potentially informative hybrid systems could not be included because of missing or incomplete parental cytogenetic records, limiting the ability to test broader hypotheses about hybrid chromosomal architecture in less-studied or recently described taxa.

These gaps suggest caution when generalizing the patterns observed in this study to all monocot lineages. Broader sampling of chromosome counts, ideally with renewed investment in cytogenetic field and laboratory work, would improve the resolution of hybrid analyses and allow for more powerful meta-analyses using the PDI framework. In particular, expanding cytogenetic datasets to include both parental taxa and hybrids from underrepresented clades or regions would greatly enhance confidence in conclusions about hybrid viability, the role of polyploidy, and patterns of chromosomal divergence. Future work that combines traditional chromosome counting with modern genomic tools could help fill these gaps and advance a more complete understanding of hybrid evolution.

### Summary

In summary, this study provides the first broad-scale, quantitative assessment of cytogenetic patterns in monocot hybridization using the ploidy deviation index (PDI), revealing meaningful relationships between hybrid chromosome configurations and factors such as hybrid origin, growth habit, genus size, and genus chromosome range. The results showed that homoploid hybrids were far more common than traditionally expected, with no clear evidence that polyploid hybrids dominate across monocot systems. Contrary to long-held assumptions, homoploid hybrids were not disproportionately limited to artificial systems, suggesting they can persist and establish in natural habitats as well.

Additionally, the study revealed significant differences across growth habits, with aquatic systems more likely to harbor homoploid hybrids than graminoids and geophytes significantly more likely to produce polyploid hybrids as compared to aquatics. This pattern reflects how stable aquatic environments may favor maintenance of parental chromosome structures while geophytes, exposed to periodic dormancy and environmental fluctuations, may benefit from polyploidy to stabilize meiosis and enhance adaptability. The results also demonstrate that intermediate hybrids are particularly common in larger, more chromosomally variable genera, while homoploid hybrids tend to arise in smaller, more karyotypically uniform lineages.

Polyploid hybrids, in contrast, displayed the highest PDI values in large and chromosomally diverse genera, supporting their role as a mechanism for stabilizing meiosis in the face of extreme parental karyotype differences. These patterns challenge traditional expectations about the predominance of polyploid hybridization over homoploid hybridization and highlight the importance of ecological and taxonomic context in shaping hybrid outcomes.

Future work should aim to apply the PDI framework across other plant lineages, including eudicots, magnoliids, and non-angiosperm lineages, to test whether these patterns are consistent more broadly throughout the plant kingdom. Incorporating genomic and ecological data to test reproductive isolation and hybrid lineage persistence would further strengthen these inferences in the context of hybrid speciation. Additionally, expanding sampling in underrepresented lineages and growth habits will help clarify whether the observed trends hold universally. Ultimately, the PDI offers a flexible and standardized tool for quantifying hybrid chromosomal relationships, providing a valuable resource for meta-analyses that integrate cytogenetics, systematics, and evolutionary biology to assess the cytogenetic constraints of hybridization.

## Supplemental Information

The supplemental information for the manuscript includes two figures and one appendix. Figure S1 presents a violin plot with an overlaid box-and-whisker plot illustrating the full distribution of Ploidy Deviation Index (PDI) scores across all hybrid cases, regardless of life history category, with ranges for uniparentally homoploid, intermediate, polyploid, and homoploid hybrids clearly distinguished. Figure S2 summarizes the temporal distribution of cytogenetic literature by showing the frequency of dataset publications containing karyotypic information across publication years, overlaid with a bell curve to visualize the trend. Appendix S1 provides the comprehensive hybrid system dataset, including ploidy information for hybrids and their parental species, associated reference citations, and metadata such as genus size, growth habit, and origin type

## Acknowledgements

This research was made possible through support from the Lewis and Clark Fund for Exploration and Research, the Torrey Botanical Society, the Society of Systematic Biologists, the Mario Einaudi Center of International Studies, and the Moore Fund of the L.H. Bailey Hortorium Herbarium. Additionally, the stipend of the first author was supported in part by the Tara Atluri Memorial Fund.

