## Supplemental Table 1 for "Cytogenetic constraints on hybridization: A meta-analysis investigating the role of chromosome number in monocot hybrid evolution using a newly developed tool, the ploidy deviation index (PDI)"

*Supplemental Information*

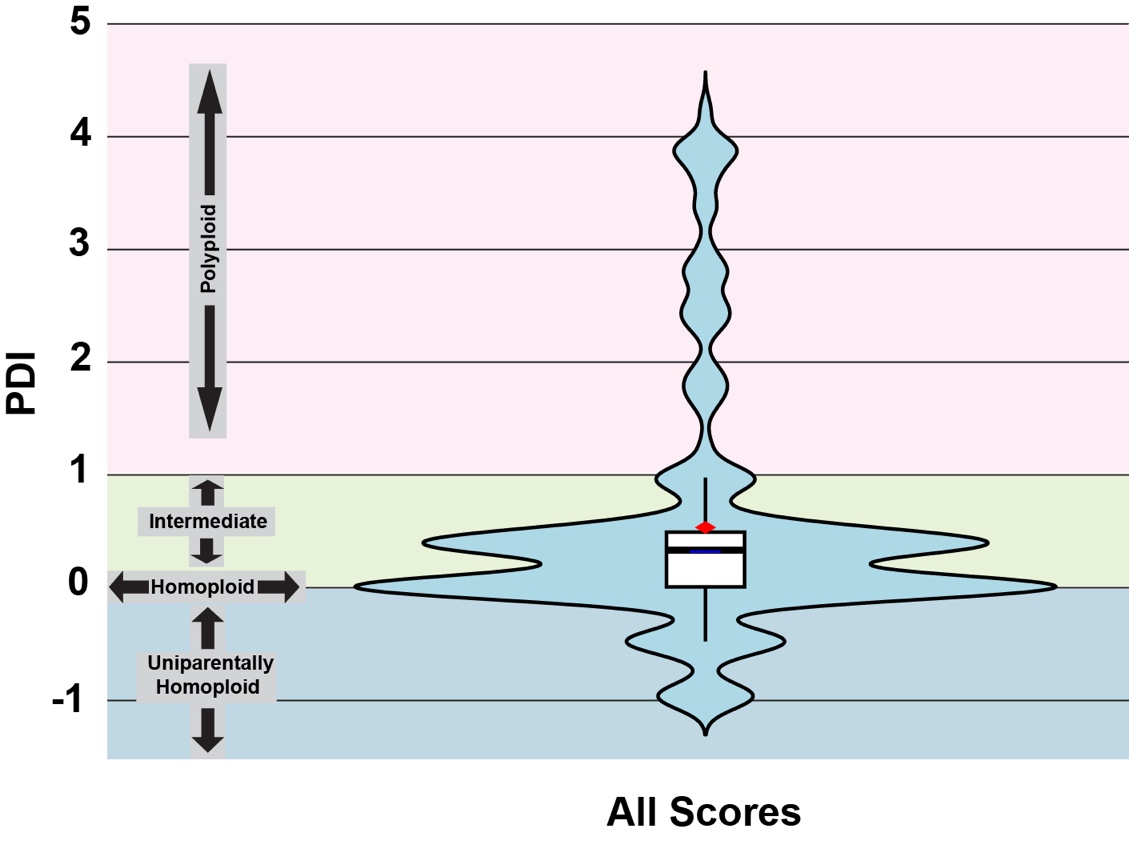

**Figure S1.** Violin plot with box and whisker plot overlay illustrating the overall distribution of PDI score regardless of life history category. Mean PDI score represented by the red diamond. Categories representing the PDI scores is indicated on left, with the range for uniparentally homoploid shaded in blue, the range for Intermediate hybrid shaded in green, and the range for polyploid shaded in pink. The PDI for homoploid hybrids is exactly 0.

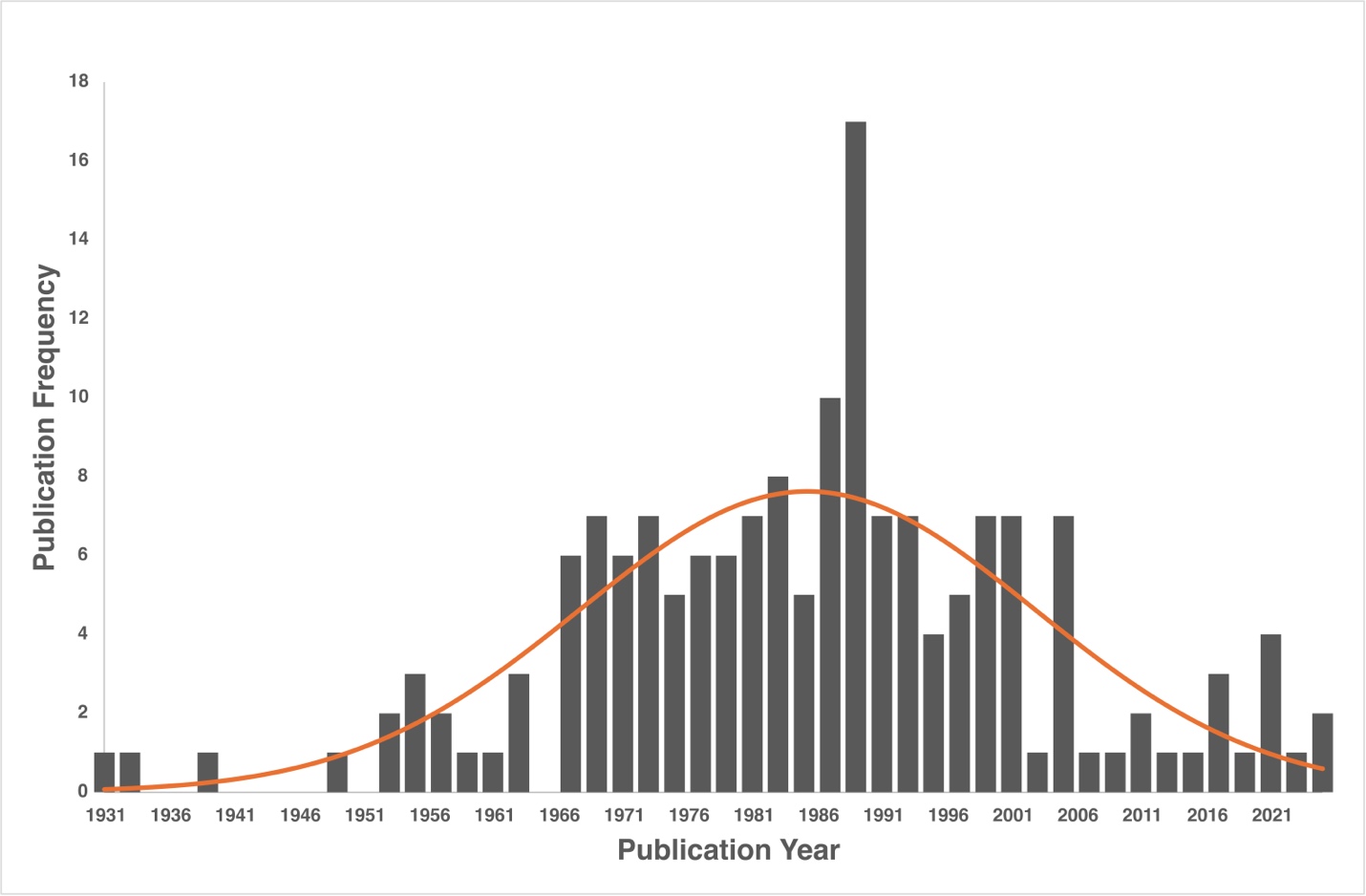

**Figure S2.** Frequency of dataset publications including karyotypic information by publication year. The orange line represents a bell curve to better visualize the data distribution. For a list of publications, see S1.

**Appendix S1.** Hybrid system dataset with ploidy information and reference citations.

| **Genus Size** | **Habit** | **Origin Type** | **Parent 2 citation** | **P2 Ploidy** | **Parent Species 2** | **Parent 1 citation** | **P1 ploidy** | **Parent Species 1** | **Hybrid citation** | **Hybrid ploidy** | **Hybrid Species** | **Family** | **ID** |
| --- | --- | --- | --- | --- | --- | --- | --- | --- | --- | --- | --- | --- | --- |
| 225 | Evergreen | natural | Pinkava and Baker 1985 | 2n = 60 | A. schottii var. schottii | Pinkava and Baker 1985 | 2n = 60 | A. deserti var. simplex (Gentry) W. C. Hodgs. and Reveal | Rice et al. 2014 | 2n = 90 | Agave × ajoensis W. C. Hodgs. | Asparagaceae | 1 |
| 225 | Evergreen | natural | Pinkava and Baker 1985 | 2n = 60 | A. toumeyana var. bella Breitung | Pinkava and Baker 1985 | 2n = 60 | A. chrysantha Peebles | Pinkava and Baker 1985 | 2n = 60 | Agave × arizonica Gentry and J. H. Weber | Asparagaceae | 2 |
| 1089 | Geophyte | natural | Liu and Zhou 1988 | 2n = 16 | A. fistulosum L. | Májovský et al. 1978 | 2n = 16 | A. cepa L. | Zakirova and Nafanailova 1988 | 2n = 16 | Allium × proliferum (Moench) Schrad. ex Willd. | Amaryllidaceae | 3 |
| 13 | Geophyte | natural | D'Emerico et al. 1996 | 2n =32 | A. papilionacea. | Rice et al. 2014 | 2n =36 | A. morio subsp. morio | D’Emerico et al. 1994 | 2n =34 | Anacamptis × gennarii (Rchb.) | Orchidaeceae | 4 |
| 1357 | Vine | artificial | Petersen 1989 | 2n = 30 | A. pedatoradiatum Schott | Sheffer and Croat 1983 | 2n = 35 | A. leuconeurum Lem. | Sharma and Bhattacharya 1966 | 2n = 30 | Anthurium × macrolobium hort. ex W. Bull | Araceae | 5 |
| 34 | Evergreen | natural | Carvalho and Bandel 1986 | 2n=32 | A. speciosa Mart. | Carvalho and Bandel 1986 | 2n=32 | A. eichleri (Drude) A. J. Hend. | Carvalho and Bandel 1986 | 2n=32 | Attalea × teixeirana (Bondar) Zona | Arecaceae | 6 |
| 62 | Epiphyte | artificial | Fedrov 1969 | 2n = 54 | Quesnelia liboniana (D. Jonghe) Mez | Ramírez-Morillo and Brown 2001 | 2n = 50 | B. nutans H. Wendl. ex Regel | Fedrov 1969 | 2n = 54 | Billbergia × perringiana Wittm. | Bromeliaceae | 7 |
| 178 | Graminoid | natural | Májovský et al. 1987 | 2n = 28 | C. epigejos (L.) Roth | Májovský et al. 1974 | 2n = 28 | C. arundinacea (L.) Roth | Sorokin 1993 | 2n = 28 | Calamagrostis × acutiflora (Schrad.) DC. | Poaceae | 8 |
| 178 | Graminoid | natural | Májovský et al. 1987 | 2n = 28 | C. epigejos (L.) Roth | Wenworth et al. 1991 | 2n = 28 | C. arenaria Roth | Westergaard 1943 | 2n = 56 | Calamagrostis × baltica | Poaceae | 9 |
| 178 | Graminoid | natural | Mičieta 1986 | 2n = 28 | C. canescens (Weber) Roth | Májovský et al. 1974 | 2n = 28 | C. arundinacea (L.) Roth | Paszko 2006 | 2n = 28 | Calamagrostis × hartmaniana Fr. | Poaceae | 10 |
| 12 | Evergreen | artificial | Hanson et al. 2001 | 2n = 18 | C. indica L. | Segeren and Mass 1971 | 2n =18 | C. glauca | Chen et al. 2003 | 2n =18 | Canna × hybrida Rodigas | Cannaceae | 11 |
| 2094 | Graminoid | natural | Lipnerová et al. 2012 | 2n = 70 | C. dacica Heuff. | Luceño and Aedo 1994 | 2n = 84 | C. nigra (L.) Reichard | Więcław et al. 2020 | 2n = 74 | Carex × decolorans Wimm. | Cyperaceae | 12 |
| 2094 | Graminoid | natural | Cayouette and Morisset 1985 | 2n = 76 | C. aquatilis Wahlenb. | Zhukova and Petrovsky 1980 | 2n=80 | C. subspathacea Wormsk. ex Hornem. | Cayouette and Morisset 1985. | 2n = 79 | Carex × flavicans (F.Nyl.) F.Nyl. | Cyperaceae | 13 |
| 2094 | Graminoid | natural | Cayouette and Morisset 1985 | 2n = 76 | C. aquatilis Wahlenb. | Cayouette and Morisset 1985 | 2n = 72 | C. paleacea Schreb. ex Wahlenb. | Cayouette and Morisset 1985 | 2n = 74 | Carex × neofilipendula Lepage | Cyperaceae | 14 |
| 2094 | Graminoid | natural | Więcław et al. 2020 | 2n = 78 | C. randalpina B.Walln. | Murín and Feráková 1986 | 2n = 84 | C. acuta L. | Więcław et al. 2020 | 2n = 84 | Carex × oenensis A. Neumann ex B. Walln. | Cyperaceae | 15 |
| 2094 | Graminoid | natural | Cayouette and Morisset 1985 | 2n = 84 | C. nigra (L.) Reichard | Zhukova and Petrovsky 1980 | 2n=80 | C. subspathacea Wormsk. ex Hornem. | Cayouette and Morisset 1985 | 2n=81 | Carex × reducta Drejer | Cyperaceae | 16 |
| 2094 | Graminoid | natural | Cayouette and Morisset 1985 | 2n = 72 | C. paleacea Schreb. ex Wahlenb. | Cayouette and Morisset 1985 | 2n = 73 | C. recta Boott | Cayouette and Morisset 1985. | 2n = 73 | Carex × saxenii Raymond | Cyperaceae | 17 |
| 2094 | Graminoid | natural | Luceño and Aedo 1994 | 2n = 84 | C. nigra (L.) Reichard | Cayouette and Morisset 1985 | 2n=77 | C. salina Wahlenb. | Cayouette and Morisset 1985. | 2n = 80 | Carex × spiculosa Fr. | Cyperaceae | 18 |
| 2094 | Graminoid | natural | Cayouette and Morisset 1985 | 2n = 72 | C. paleacea Schreb. ex Wahlenb. | Cayouette and Morisset 1985 | 2n=77 | C. salina Wahlenb. | Cayouette and Morisset 1985. | 2n=74 | Carex × subpaleacea J.Cay. | Cyperaceae | 19 |
| 2094 | Graminoid | natural | Löve 1972 | 2n = 112 | C. hirta L. | Löve 1981 | 2n = 74 | C. atherodes Spreng. | Więcław et al. 2020 | 2n = 108 | Carex × walasii Ceyn.-Gield | Cyperaceae | 20 |
| 2094 | Graminoid | natural | Love and Love 1981 | 2n=72 | C. paleacea Schreb. ex Wahlenb. | Faulkner 1972 | 2n=76 | C. aquatilis Wahlenb. | Cayouette and Morisset 1985 | 2n =74 | Carex aquatilis x palaecea | Cyperaceae | 21 |
| 2094 | Graminoid | natural | Cayouette and Morsset 1985 | 2n = 73 | C. recta Boott | Faulkner 1972 | 2n=76 | C. aquatilis Wahlenb. | Cayouette and Morisset 1985 | 2n =74 | Carex aquatilis x recta | Cyperaceae | 22 |
| 2094 | Graminoid | natural | Cayouette and Morsset 1985 | 2n = 83 | C. subspathacea Wormsk. ex Hornem | Cayouette and Morisset 1985 | 2n =78 | C. aquatilis Wahlenb. | Cayouette and Morisset 1985 | 2n = 79 | *Carex aquatilis* x *subspathacea* | Cyperaceae | 23 |
| 2094 | Graminoid | natural | Love and Love 1981 | 2n=72 | C. paleacea Schreb. ex Wahlenb. | Májovský et al. 2000 | 2n=84 | C. nigra (L.) Reichard | Cayouette and Morisset 1985 | 2n = 79 | Carex nigra x palaecea | Cyperaceae | 24 |
| 2094 | Graminoid | natural | Cayouette and Morsset 1985 | 2n=73 | C. recta Boott | Májovský et al. 2000 | 2n=84 | C. nigra (L.) Reichard | Cayouette and Morisset 1985 | 2n =78 | Carex nigra x recta | Cyperaceae | 25 |
| 2094 | Graminoid | natural | Cayouette and Morsset 1985 | 2n =77 | C. salina Wahlenb. | Cayouette and Morisset 1985 | 2n = 83 | C. nigra (L.) Reichard | Cayouette and Morisset 1985 | 2n =80 | Carex nigra x salina | Cyperaceae | 26 |
| 2094 | Graminoid | natural | Zhukova and Petrovsky 1980 | 2n=80 | C. subspathacea Wormsk. ex Hornem | Cayouette and Morisset 1985 | 2n = 83 | C. nigra (L.) Reichard | Cayouette and Morisset 1985 | 2n =81 | *Carex nigra* X *subspathacea* | Cyperaceae | 27 |
| 2094 | Graminoid | natural | Cayouette and Morsset 1985 | 2n=73 | C. recta Boott | Love and Love 1981 | 2n=72 | C. paleacea Schreb. ex Wahlenb. | Cayouette and Morisset 1985 | 2n =73 | Carex paleacea x recta | Cyperaceae | 28 |
| 2094 | Graminoid | natural | Cayouette and Morsset 1985 | 2n=77 | C. salina Wahlenb. | Love and Love 1981 | 2n=72 | C. paleacea Schreb. ex Wahlenb. | Cayouette and Morisset 1985 | 2n =75 | Carex paleacea X salina | Cyperaceae | 29 |
| 205 | Epiphyte | natural | Jones and Daker 1968 | 2n =108 | Catasetum pileatum | Jones and Daker 1968 | 2n =54 | C. macrocarpum | Jones et al. 1968 | 2n =54 | Catasetum × tapiriceps Rchb. | Orchidaeceae | 30 |
| 6 | Geophyte | artificial | Sveshnikova and Zemskova 1988 | 2n = 22 | C. miniata Regel | Sveshnikova and Zemskova 1988 | 2n = 22 | C. nobilis Lindl. | Fedrov 1969 | 2n = 18 | Clivia × cyrtanthiflora (Lindl.) T. Moore | Amaryllidaceae | 31 |
| 600 | Epiphyte | artificial | Hartati et al. 2017 | 4n =72 | C. rumphii Lindl. | Hartati et al. 2017 | 2n =36 | C. pandurata Lindl. | Hartati et al. 2017 | 2n =54 | Coelogyne pandurata x Coelogyne rumphii | Orchidaceae | 32 |
| 163 | Geophyte | artificial | D'Amato 1955 | 2n = 38 | C. autumnale L. | Persson 1999 | 2n = 54 | C. cilicicum (Boiss.) Dammer | Persson 2007 | 2n = 46 | Colchicum × byzantinum Ker Gawl | Colchicaceae | 33 |
| 22 | Evergreen | natural | Röser 1993 | 2n=36 | C. macroglossa H. Wendl. ex Becc. | Dahlgren and Glassman 1963 | 2n=36 | C. hospita Mart. | Dahlgren and Glassman 1963 | 2n=36 | Copernicia × burretiana León | Arecaceae | 34 |
| 116 | Geophyte | natural | Patwary and Zaman 1981 | 2n = 22 | C. zeylanicum (L.) L. | Sinha and Roy 1986 | 2n = 22 | C. asiaticum L. | Jee and Vijayavalli 1997 | 2n = 22 | Crinum × amabile Donn ex K. Gawl. | Amaryllidaceae | 35 |
| 116 | Geophyte | artificial | Patwary and Zaman 1981 | 2n = 22 | C. zeylanicum (L.) L. | Sveshnikova and Zemskova 1988 | 2n = 22 | C. americanum L. | Fedrov 1969 | 2n = 22 | Crinum × grandiflorum auct. | Amaryllidaceae | 36 |
| 116 | Geophyte | artificial | Sinha and Roy 1986 | 2n = 22 | C. moorei Hook. | Sveshnikova and Zemskova 1988 | 2n = 22 | C. bulbispermum (Burm.) Milne-Redh. and Schweick. | Raina and Khoshoo 1971 | 2n = 22 | Crinum × powellii Baker | Amaryllidaceae | 37 |
| 8 | Geophyte | natural | Shibata 1962 | 2n = 22 | C. aurea (Pappe) Planch | Goldblatt 1971 | 2n = 22 | C. pottsii Baker | Goldblatt 1971 | 2n = 22 | Crocosmia ×crocosmiiflora (Lemoine) N.E.Br. | Iridaceae | 38 |
| 250 | Geophyte | artificual | Heywood 1983 | 2n = 8 | C. flavus Weston | Zahar'eva and Makusenko 1969 | 2n = 12 | C. angustifolius Weston | Mather 1932 | 2n = 8 | Crocus × stellaris Haw. | Iridaceae | 39 |
| 77 | Aquatic | natural | Arends et al. 1982 | 2n = 34 | C. griffithii Schott | Arends et al. 1982 | 2n = 34 | C. cordata Griff. | Jacobsen 1977 | 2n = 34 | Cryptocoryne × purpurea Ridl. | Araceae | 40 |
| 77 | Aquatic | natural | Arends et al. 1982 | 2n = 34 | C. nurii Furtado | Arends et al. 1982 | 4n = 68 | C. cordata Griff. | Bastmeijer 2001 | 3n = 54 | Cryptocoryne × timahensis Bastm. | Araceae | 41 |
| 77 | Aquatic | natural | Jacobsen 1976 | 2n = 28 | C. walkeri Schott | Jacobsen 1976 | 2n = 28 | C. parva de Wit | Jacobsen 1976 | 2n = 28 | Cryptocoryne × willisii Reitz | Araceae | 42 |
| 15 | Graminoid | natural | Hurcombe 1947 | 2n = 20 | C. transvaalensis B. Davy | Májovský et al. 1978 | 2n = 40 | C. dactylon (L.) Pers. | Darlington and Wylie 1955 | 2n = 30 | Cynodon × magennisii Hurcombe | Poaceae | 43 |
| 34 | Geophyte | natural | Májovský et al. 1978 | 2n= 80 | D. majalis (Rchb.) P. F. Hunt and Summerh | Májovský et al. 2000 | 2n=40 | D. incarnata (L.) Soó | Averyanov 1986 | 2n=60 | Dactylorhiza × ishorica Aver. | Orchidaceae | 44 |
| 1600 | Epiphyte | natural | Jones et al. 1982 | 2n =38 | D. speciosum Sm. | Solovjova 2003 | 2n =40 | D. kingianum Bidwill ex Lindl | Rice et al. 2014 | 57 | Dendrobium × delicatum F.M Bailey | Orchidaeceae | 45 |
| 24 | Geophyte | Natural | Probatova and Sokolovskaya 1981 | 2n = 16 | D. smilacinum | Lee 1979 | 2n = 16 | D. sessile | Kogi Suzuki 1988 | 2n = 16 | Disporum × hishiyamanum K. Suzuki | Colchicaceae | 46 |
| 1882 | Epiphyte | natural | Arida et al. 2021 | 6n =156 | E. orchidiflorum Salzm. ex Lindl. | Arida et al. 2021 | 2n = 52 | E. denticulatum Barb.Rodr. | Arida et al. 2021 | 4n=102 | Epidendrum x purpureum Barb.Rodr. | Orchidaeceae | 47 |
| 16 | Graminoid | natural | Dalgaard 1988 | 2n = 58 | E. scheuchzeri Hoppe | Löve 1981 | 2n = 58 | E. chamissonis C. A. Mey. | Knaben and Engelskjon 1967 | 2n = 58 | Eriophorum × medium Andersson | Cyperaceae | 48 |
| 226 | Geophyte | natural | Peruzzi and Aquaro 2005 | 2n = 36 | G. granatelli | Peruzzi 2008 | 2n = 48 | G. bohemica | Peruzzi and Bartolucci 2006 | 2n = 36 | Gagea × luberonensis J. -. Tison | Liliaceae | 49 |
| 23 | Geophyte | natural | Sveshnikova 1988 | 2n = 24 | G. woronowii Losinsk. | Sveshnikova and Zemskova 1988 | 2n =24 | G. alpinus Sosn. | Sveshnikova 1967 | 2n = 24 | Galanthus × allenii Baker | Amaryllidaceae | 50 |
| 296 | Geophyte | artificial | Goldblatt et al. 1997 | 2n = 30 | G. oppositiflorus Herb | Goldblatt 1989 | 2n = 30 | G. dalenii V Geel | Cheng et al. 1991 | 4n = 60 | Gladiolus × gandavensis V. Houtte | Iridaceae | 51 |
| 143 | Evergreen | natural | Lim 1972 | 2n=48 | G. patens Miq | Lim 1972 | 2n=48 | G. cernua Baker | Siew-Ngo 1972 | 2n=48 | Globba × intermedia S. N. Lim | Zingiberaceae | 52 |
| 42 | Graminoid | natural | Feráková and Javorčíková 1974 | 2n = 40 | G. notata Chevall. | Májovský et al. 1974 | 2n = 40 | G. fluitans (L.) R. Br. | Cave 1958 | 2n = 40 | Glyceria × pedicellata F. Towns. | Poaceae | 53 |
| 42 | Graminoid | natural | Masumura 1989 | 2n = 20 | G. leptolepis Ohwi | Masumura 1989 | 2n = 40 | G. ischyroneura Steud. | Masumura 1989 | 2n = 30 | Glyceria × tokitana Masamura | Poaceae | 54 |
| 28 | Geophyte | natural | Teppner and Klein 1990 | 2n =40 | G. rhellicani. | Marhold et al. 2005 | 2n =40 | G. conopsea  (L.) R.Br. | Eccarius 2022 | 3n = 54 | Gymnadenia × suaveolens (Vill.) Rchb.f. | Orchidaeceae | 55 |
| 31 | Graminoid | natural | Sauer 1970 | 2n = 14 | H. petzense H. Melzer | Gervais 1966 | 2n = 14 | H. parlatorei  (J. Woods) Pilg. | Sauer 1970 | 2n = 14 | Helictotrichon × krischae H. Melzer | Poaceae | 56 |
| 12 | Geophyte | natural | Castroviejo and Feliner 1986 | 2n = 16 | H. hispanica (Mill.) Rothm. | Martinez 1997 | 2n = 16 | H. non-scripta (L.) Chouard ex Rothm. | Vallejo-Marín and Hiscock 2016 | 2n = 16 | Hyacinthoides × massartiana Geerinck | Asparagaceae | 57 |
| 12 | Geophyte | natural | Castroviejo and Feliner 1986 | 2n = 24 | H. hispanica (Mill.) Rothm. | Martinez 1997 | 2n = 24 | H. non-scripta (L.) Chouard ex Rothm. | Vallejo-Marín and Hiscock 2016 | 2n = 24 | Hyacinthoides × massartiana Geerinck | Asparagaceae | 58 |
| 67 | Geophyte | artificial | Fedorov 1969 | 2n = 46 | H. speciosa (L.) Salisb. | Zemskova and Sveshnikova 1999 | 2n = 44 | H. narcissiflora (Jacq.) J. F. Macbr. | Fedorov 1969 | 2n = 46 | Hymenocallis × macrostephana Baker | Amaryllidaceae | 59 |
| 67 | Geophyte | artificial | Fedorov 1969 | 2n = 46 | H. speciosa (L.) Salisb. | Zemskova and Sveshnikova 1999 | 2n = 44 | H. narcissiflora (Jacq.) J. F. Macbr. | Fedrov 1969 | 2n = 96 | Hymenocallis × macrostephana Baker | Amaryllidaceae | 60 |
| 316 | Geophyte | natural | Májovský et al. 1978 | 2n = 24 | I. variegata L. | Mitra 1956 | 2n = 24 | I. pallida Lam. | Colasante and Ricci 1975 | 2n = 44 | Iris × germanica L. | Iridaceae | 61 |
| 122 | Geophyte | aritificial | Kogi and Ogihara 1988 | 2n = 24 | L. maculatum | Bambacioni-Mezzetti 1931 | 2n = 24 | L. bulbuferum | Rice et al. 2014 | 2n = 24 | Lilium × hollandicum Bergmans | Liliaceae | 62 |
| 25 | Geophyte | natural | Tae and Ko 1987 | 2n = 22 | L. sanguinea var. koreana (Nakai) T. Koyama | Chen and Li 1985 | 2n = 16 | L. chinensis Traub | Tae and Ko 1993 | 2n = 30 | Lycoris × chejuensis K. H. Tae and S. C. Ko | Amaryllidaceae | 63 |
| 40 | Geophyte | natural | Nishikawa 1979 | 2n=36 | M. dilatatum (A. Wood) A. Nelson and J. F. Macbr. | Májovský et al. 1970 | 2n = 36 | M. bifolium (L.) F. W. Schmidt | Probatova and Sokolovskaya 1986 | 2n = 60 | Maianthemum × intermedium Vorosch. | Asparagaceae | 64 |
| 14 | Graminoid | natural | Nishiwaki et al. 2011 | 2n =38 | M. sinensis Andersson | Fedorov 1969 | 2n = 64 | M. sacchariflorus (Maxim.) Hack. | Hodkinson and Renvoize 2001 | 2n = 57 | *Miscanthus* × *longiberbis* (Hack.) Nakai | Poaceae | 65 |
| 85 | Evergreen | natural | Wang et al. 1994 | 2n=22 | M. balbisiana Colla | Wang et al. 1994 | 2n=22 | M. acuminata Colla | Cheesman and Larter 1935 | 2n=22 | Musa × paradisiaca L. | Musaceae | 66 |
| 76 | Geophyte | artificial | Ruz and Saudo 1976 | 2n = 14 | N. cuatrecasasii Fern.Casas, M.Laínz and Ruíz Rejón | Wylie 1952 | 2n = 14 | N. atlanticus Stern | Brandham and Kirton 1987 | 2n = 14 | N. atlanticus × N. asturiensis | Amaryllidaceae | 67 |
| 76 | Geophyte | artificial | Alcaraz 1983 | 2n = 14 | N. gaditanus Boiss. and Reut. | Darlington and Wylie 1955 | 2n = 14 | N. rupicola Dufour | Brandham and Kirton 1987 | 2n = 14 | N. rupicola × N. gaditanus | Amaryllidaceae | 68 |
| 76 | Geophyte | artificial | Maugini 1953 | 2n = 20 | N. tazetta 'Soleil d'Or' | Darlington and Wylie 1955 | 2n = 14 | N. cyclamineus DC. | Brandham and Kirton 1987 | 2n = 17 | Narcissus 'Cyclataz' | Amaryllidaceae | 69 |
| 76 | Geophyte | artificial | Maugini 1953 | 2n = 20 | N. tazetta L. | Cesca 1972 | 2n = 14 | N. poeticus L. | Brandham and Kirton 1987 | 2n = 17 | Narcissus × biflorus Curtis | Amaryllidaceae | 70 |
| 76 | Geophyte | natural | Darlington and Wylie 1955 | 2n = 14 | N. jonquilla L. | Maugini 1953 | 2n = 20 | N. tazetta L. | Darlington and Wylie 1955 | 2n = 17 | Narcissus × compressus Haw. | Amaryllidaceae | 71 |
| 76 | Geophyte | artificial | Maugini 1953 | 2n = 20 | N. tazetta L. | Darlington and Wylie 1955 | 2n = 14 | N. jonquilla L. | Brandham and Kirton 1987 | 2n = 31 | Narcissus × compressus Haw. | Amaryllidaceae | 72 |
| 76 | Geophyte | artificial | Darlington and Wylie 1955 | 2n = 14 | N. cyclamineus DC. | Maugini 1953 | 2n=22 | N. tazetta L. | Rice et al. 2014 | 2n = 17 | Narcissus × cyclazetta Chater and Stace | Amaryllidaceae | 73 |
| 76 | Geophyte | artificial | Alcaraz 1983 | 2n = 14 | N. juncifolius Req. ex Lag. | D'Amato et al. 2004 | 2n = 22 | N. papyraceus Ker Gawl. | Darlingtong and Wylie 1955 | 2n = 50 | Narcissus × dubius Gouan | Amaryllidaceae | 74 |
| 76 | Geophyte | natural | Barra and González 1984 | 2n = 14 | N. hispanicus Gouan | Montserrat and Vives 1991 | 2n = 14 | N. triandrus L. | Fernandes 1977 | 2n = 14 | Narcissus × hannibalis A. Fern. | Amaryllidaceae | 75 |
| 76 | Geophyte | artificial | Baldini 1993 | 2n = 14 | N. pseudonarcissus L. | Cesca 1972 | 2n = 14 | N. poeticus L. | Brandham and Kirton 1987 | 2n = 14 | Narcissus × incomparabilis Mill. | Amaryllidaceae | 76 |
| 76 | Geophyte | natural | Baldini 1993 | 2n = 14 | N. pseudonarcissus L. | Cesca 1972 | 2n = 14 | N. poeticus L. | Heitz 1926 | 2n=14 | Narcissus × incomparabilis Mill. | Amaryllidaceae | 77 |
| 76 | Geophyte | artificial | Maugini 1953 | 2n = 20 | N. tazetta L. | Darlington and Wylie 1955 | 2n = 14 | N. jonquilla L. | Brandham and Kirton 1987 | 2n = 17 | Narcissus × intermedius Loisel | Amaryllidaceae | 78 |
| 76 | Geophyte | natural | Cesca 1972 | 2n = 14 | N. poeticus L. | Maugini 1953 | 2n=22 | N. tazetta L. | Rice et al. 2014 | 2n = 17 | Narcissus × medioluteus Mill. | Amaryllidaceae | 79 |
| 76 | Geophyte | natural | Darlington and Wylie 1955 | 2n = 14 | N. cyclamineus DC. | Baldini 1993 | 2n = 14 | N. pseudonarcissus L. | Rice et al. 2014 | 3n = 21 | Narcissus × monochromus P. D. Sell | Amaryllidaceae | 80 |
| 76 | Geophyte | artificial | Baldini 1993 | 2n = 14 | N. pseudonarcissus L. | Darlington and Wylie 1955 | 2n = 14 | N. jonquilla L. | Brandham and Kirton 1987 | 2n = 14 | Narcissus × odorus L. var plenus Hort | Amaryllidaceae | 81 |
| 76 | Geophyte | artificial | Baldini 1993 | 2n = 14 | N. pseudonarcissus L. | Darlington and Wylie 1955 | 2n = 14 | N. jonquilla L. | Brandham and Kirton 1987 | 2n = 14 | Narcissus × odorus L. var rugulosus Hort | Amaryllidaceae | 82 |
| 76 | Geophyte | artificial | Baldini 1993 | 2n = 14 | N. pseudonarcissus L. | Darlington and Wylie 1955 | 2n = 14 | N. jonquilla L. | Brandham and Kirton 1987 | 2n = 21 | Narcissus × odorus L. var rugulosus Hort | Amaryllidaceae | 83 |
| 76 | Geophyte | natural | D'Amato et al. 2004 | 2n = 30 | N. serotinus L. | Darlington $ Wylie 1955 | 2n = 20 | N. elegans (Haw.) Spach | Fernandes 1967 | 2n = 20 | Narcissus × perangustus F. Casas | Amaryllidaceae | 84 |
| 76 | Geophyte | natural | Montserrat and Vives 1991 | 2n = 14 | N. assoanus Dufour ex Schult. and Schult. | Darlington and Wylie 1955 | 2n = 50 | N. dubius Gouan | Fernández 1983 | 2n = 32 | Narcissus × pujolii F. Quer | Amaryllidaceae | 85 |
| 76 | Geophyte | natural | Fernandes 1968 | 2n = 14 | N. portensis Pugsley | Montserrat and Vives 1991 | 2n = 14 | N. triandrus L. | Darlington and Wylie 1955 | 2n = 21 | Narcissus × taitii Henriq. | Amaryllidaceae | 86 |
| 76 | Geophyte | artificial | Galland 1988 | 2n = 14 | N. watieri Maire | Darlington and Wylie 1955 | 2n = 14 | N. calcicola Mendonça | Brandham and Kirton 1987 | 2n = 14 | Narcissus calcicola × Narcissus watieri | Amaryllidaceae | 87 |
| 76 | Geophyte | artificial | Maugini 1953 | 2n = 20 | N. tazetta L. | Cesca 1972 | 2n = 14 | N. poeticus L. | Brandham and Kirton 1987 | 2n = 17 | Narcissus cv. ‘Alsace’ | Amaryllidaceae | 88 |
| 76 | Geophyte | artificial | Maugini 1953 | 2n = 20 | N. tazetta L. | Cesca 1972 | 2n = 14 | N. poeticus L. | Brandham and Kirton 1987 | 2n = 17 | Narcissus cv. ‘Cragford’ | Amaryllidaceae | 89 |
| 76 | Geophyte | artificial | Brandham and Kirton 1987 | 2n = 24 | N. tazetta L. | Brandham and Kirton 1987 | 2n = 28 | N. poeticus L. | Brandham and Kirton 1987 | 2n = 24 | Narcissus cv. ‘Elvira’ | Amaryllidaceae | 90 |
| 76 | Geophyte | artificial | Maugini 1953 | 2n = 20 | N. tazetta L. | Cesca 1972 | 2n = 14 | N. poeticus L. | Brandham and Kirton 1987 | 2n = 17 | Narcissus cv. ‘Geranium’ | Amaryllidaceae | 91 |
| 76 | Geophyte | artificial | Brandham and Kirton 1987 | 2n = 24 | N. tazetta L. | Brandham and Kirton 1987 | 2n = 28 | N. poeticus L. | Brandham and Kirton 1987 | 2n = 24 | Narcissus cv. ‘Glorious’ | Amaryllidaceae | 92 |
| 76 | Geophyte | artificial | Maugini 1953 | 2n = 22 | N. tazetta L. | Rice et al. 2015 | 2n = 28 | N. poeticus L. | Brandham and Kirton 1987 | 2n = 24 | Narcissus cv. ‘Golden Perfection’ | Amaryllidaceae | 93 |
| 76 | Geophyte | artificial | Maugini 1953 | 2n = 24 | N. cv. ‘Golden Perfection’ | Darlington and Wylie 1955 | 2n = 14 | N. jonquilla L. | Brandham and Kirton 1987 | 2n = 31 | Narcissus cv. ‘Golden Perfection’ | Amaryllidaceae | 94 |
| 76 | Geophyte | artificial | Brandham and Kirton 1987 | 2n = 22 | N. tazetta hermione (Haw.) Spach | Brandham and Kirton 1987 | 2n = 20 | N. tazetta hermione (Haw.) Spach | Brandham and Kirton 1987 | 2n = 31 | Narcissus cv. ‘Highfield Beauty’ | Amaryllidaceae | 95 |
| 76 | Geophyte | artificial | Barra and Gonzalez 1984 | 2n = 14 | N.pseudonarcissus subsp. bicolor (L.) Baker | Rice et al. 2014 | 2n = 21 | N. cv 'Maximus' (N. pseudonarcissus subsp. major Baker) | Brandham and Kirton 1987 | 2n = 21/22 | Narcissus cv. ‘Horsefieldii’ | Amaryllidaceae | 96 |
| 76 | Geophyte | artificial | Maugini 1953 | 2n = 20 | N. tazetta L. | Cesca 1972 | 2n = 14 | N. poeticus L. | Brandham and Kirton 1987 | 2n = 17 | Narcissus cv. ‘Irene’ | Amaryllidaceae | 97 |
| 76 | Geophyte | artificial | Maugini 1953 | 2n = 22 | N. tazetta L. | Cesca 1972 | 2n = 14 | N. poeticus L. | Brandham and Kirton 1987 | 2n = 17 | Narcissus cv. 'Jaune a Merveille’ | Amaryllidaceae | 98 |
| 76 | Geophyte | artificial | Maugini 1953 | 2n = 22 | N. tazetta L. | Cesca 1972 | 2n = 14 | N. poeticus L. | Brandham and Kirton 1987 | 2n = 17 | Narcissus cv. 'L'Innocence’ | Amaryllidaceae | 99 |
| 76 | Geophyte | artificial | Maugini 1953 | 2n = 22 | N. tazetta L. | Cesca 1972 | 2n = 14 | N. poeticus L. | Brandham and Kirton 1987 | 2n = 17 | Narcissus cv. 'Laurens Koster’ | Amaryllidaceae | 100 |
| 76 | Geophyte | artificial | Daz et al. 1990 | 2n = 14 | N. asturiensis (Jord.) Pugsley | Wylie 1952 | 2n = 14 | N. atlanticus Stern | Brandham and Kirton 1987 | 2n = 14 | Narcissus cv. ‘Navarre’ | Amaryllidaceae | 101 |
| 76 | Geophyte | artificial | Daz et al. 1990 | 2n = 14 | N. asturiensis (Jord.) Pugsley | Darlington and Wylie 1955 | 2n = 14 | N. rupicola Dufour | Brandham and Kirton 1987 | 2n = 14 | Narcissus cv. ‘Navarre’ | Amaryllidaceae | 102 |
| 76 | Geophyte | artificial | Maugini 1953 | 2n = 22 | N. tazetta L. | Cesca 1972 | 2n = 14 | N. poeticus L. | Brandham and Kirton 1987 | 2n = 17 | Narcissus cv. 'Scarlet Gem’ | Amaryllidaceae | 103 |
| 76 | Geophyte | artificial | Brandham and Kirton 1987 | 2n = 22 | N. tazetta hermione (Haw.) Spach | Brandham and Kirton 1987 | 2n = 20 | N. tazetta hermione (Haw.) Spach | Brandham and Kirton 1987 | 2n = 32 | Narcissus cv. ‘Scilly White’ | Amaryllidaceae | 104 |
| 76 | Geophyte | artificial | Maugini 1953 | 2n = 22 | N. tazetta L. | Rice et al. 2014 | 2n = 14 | N. poeticus L. | Brandham and Kirton 1987 | 2n = 17 | Narcissus cv. ‘Sir Winston Churchill ’ | Amaryllidaceae | 105 |
| 76 | Geophyte | artificial | Maugini 1953 | 2n = 22 | N. tazetta L. | Cesca 1972 | 2n = 14 | N. poeticus L. | Brandham and Kirton 1987 | 2n = 17 | Narcissus cv. 'St Agnes’ | Amaryllidaceae | 106 |
| 76 | Geophyte | artificial | Galland 1988 | 2n = 14 | N. watieri Maire | Alcaraz 1983 | 2n = 14 | N. gaditanus Boiss. and Reut. | Brandham and Kirton 1987 | 2n = 14 | Narcissus gaditanus × Narcissus watieri | Amaryllidaceae | 107 |
| 76 | Geophyte | artificial | Galland 1988 | 2n = 14 | N. watieri Maire | Darlington and Wylie 1955 | 2n = 14 | N. jonquilla L. | Brandham and Kirton 1987 | 2n = 14 | Narcissus jonquilla × Narcissus watieri | Amaryllidaceae | 108 |
| 76 | Geophyte | artificial | Galland 1988 | 2n = 14 | N. watieri Maire | Darlington and Wylie 1955 | 2n = 14 | N. scaberulus Henriq. | Brandham and Kirton 1987 | 2n = 14 | Narcissus scaberulus × Narcissus watieri | Amaryllidaceae | 109 |
| 91 | Geophyte | natural | Ravenna 1991 | 2n = 19 | N. entrerianum Ravenna | Tanaka and Ohta 1982 | 2n = 19 | N. gracile (Aiton) Stearn | Rice et al. 2014 | 2n = 19 | Nothoscordum × borbonicum Kunth | Amaryllidaceae | 110 |
| 335 | Epiphyte | natural | Phang et al. 1979 | 2n=42 | T. variegata (Sw.) Braem | Braem 1988 | 2n=40 | T. haitiensis (Leonard and Ames) Braem | Rice et al. 2014 | 2n=42 | Oncidium × ann-hadderae Moir | Orchidaceae | 111 |
| 335 | Epiphyte | artificial? | Charanasri and Kamemoto 1975 | 2n =42 | T. leiboldii (Rchb.f.) Braem | Braem 1988 | 2n =42 | T. variegata (Sw.) Braem | Charanasri and Kamemoto 1975 | 2n =42 | Oncidium × cubense Moir | Orchidaeceae | 112 |
| 25 | Geophyte |  | Turco et al. 2024 | 2n = 36 | O. apulica O. Danesch and E. Danesch | Turco et al. 2024 | 2n = 36 | O. incubacea Bianca | Bianco et al. 1988 | 2n = 36 | Ophrys × franciniae Bianco and al. | Orchidaeceae | 113 |
| 25 | Geophyte | natural | D'Emerico et al. 2005 | 2n =36 | O. tenthredinifera. Willd | Bianco et al. 1988 | 2n = 36 | O. holosericea subsp. candica (E. Nelson) Renz and Taubenheim | Corrias 1983 | 2n = 36 | Ophrys × maremmae O. Danesch and E. Danesch | Orchidaeceae | 114 |
| 25 | Geophyte | natural | Druskovic and Lovka 1995 | 2n = 36 | O. sphegodes Mill. | D'Emerico et al. 2005 | 2n = 36 | O. bertolonii Moretti | Bianco et al. 1991 | 2n = 38 | Ophrys × monopolitana H. Baumann and Künkele | Orchidaeceae | 115 |
| 25 | Geophyte | natural | Bianco et al. 1988 | 2n = 36 | O. holosericea subsp. candica (E. Nelson) Renz and Taubenheim | D'Emerico et al. 2005 | 2n = 36 | O. bombyliflora (syn Ophrys fuciflora (F. W. Schmidt) Moench) | Bianco et al. 1991 | 2n = 36 | Ophrys × resurrecta O. Danesch and E. Danesch | Orchidaeceae | 116 |
| 25 | Geophyte | natural | D'Emerico et al. 2005 | 2n =36 | O. tenthredinifera. Willd | D'Emerico et al. 2005 | 2n = 36 | O. bombyliflora (syn Ophrys fuciflora (F. W. Schmidt) Moench) | Corrias 1983 | 2n = 36 | Ophrys × sommieri E. G. Camus ex Cortesi | Orchidaeceae | 117 |
| 25 | Geophyte | natural | D'Emerico et al. 2005 | 2n =36 | O. tenthredinifera. Willd | Bianco et al. 1988 | 2n =36 | O. holosericea subsp. candica | D'Emerico et al. 2005 | 2n =36 | Ophrys × tardans O. Danesch and E. Danesch | Orchidaeceae | 118 |
| 25 | Geophyte | natural | Druskovic and Lovka 1995 | 2n = 36 | O. sphegodes Mill. | Scrugli 1977 | 2n =36 | O. holosericea (Burm.f.) Greuter | Scrugli 1977 | 2n =36 | Ophrys ×arachnitiformis Gren. and M.Philippe | Orchidaeceae | 119 |
| 25 | Geophyte | natural | Hautzinger 1978 | 2n =80 | O. patens subsp. patens | Del Prete 1977 | 2n =42 | O. mascula | Hautzinger 1978 | 2n =42 | Orchis × ligustica nothosubsp. ligustica | Orchidaeceae | 120 |
| 82 | Epiphyte | artificial | Lin et al. 2001 | 3n =57 | P. aphrodite Rchb.f. | Lee et al. 2020 | 4n =76 | P. amabilis (L.) Blume | Lee et al. 2020 | 3n =57 | Phalaenopsis "Tropic Snowball" | Orchidaceae | 121 |
| 82 | Epiphyte | artificial | Kao et al. 2001 | 2n =38 | P. parishii Rchb.f | Lin et al. 2001 | 2n =38 | P. pulcherrima (Lindl.) J.J.Sm. | Lee et al. 2020 | 3n =57 | Phalaenopsis Liu's Berry 'SW' | Orchidaceae | 122 |
| 154 | Geophyte | natural | Kapoor et al. 1987 | 2n =42 | P. psycodes (L.) Lindl. | Bent 1969 | 2n =42 | P. lacera | Bent 1969 | 2n =42 | Platanthera × andrewsii | Orchidaeceae | 123 |
| 582 | Graminoid | natural | Rice et al. 2014 | 2n=42 | P. pratensis L. | Contandriopoulo and Gamisans 1974 | 2n=32 | P. alpina L. | Darlington 1955 | 2n=64 | Poa × herjedalica H. Sm. | Poaceae | 124 |
| 582 | Graminoid | natural | Engelskjøn 1979 | 2n=42 | P. flexuosa Sm. | Contandriopoulo and Gamisans 1974 | 2n=32 | P. alpina L. | Love 1948 | 2n=37 | Poa × jemtlandica (Almq.) K. Richt. | Poaceae | 125 |
| 582 | Graminoid | natural | Hernndez 1979 | 2n =14 | P. supina | Pogan et al. 1982 | 2n =28 | P. annua L. | Hand and Gregor 2011 | 2n =21 | Poa × nannfeldtii V. Jirásek | Poaceae | 126 |
| 582 | Graminoid | natural | Skalinska 1954 | 2n=64 | P. granitica Braun-Blanq. | Contandriopoulo and Gamisans 1974 | 2n=32 | P. alpina L. | Skalińska et al. 1957 | 2n=72 | Poa × nobilis Skalinska | Poaceae | 127 |
| 582 | Graminoid | natural | Murray et al. 2005 | 2n=28 | P. foliosa. (Hook) Hook | Murray et al. 2005 | 2n=28 | P. astonii Petrie | Hair 1968 | 2n=42 | Poa × poppelwellii Petrie | Poaceae | 128 |
| 79 | Geophyte | natural | Wang et al. 1987 | 2n = 18 | P. involucratum (Franch. and Sav.) Maxim. | Wang et al. 1987 | 2n = 20 | P. humile Fisch. ex Maxim. | Han et al. 1998 | 2n = 18 | Polygonatum × desoulavyi Kom. | Asparagaceae | 129 |
| 79 | Geophyte | natural | Májovský et al. 1970 | 2n = 20 | P. odoratum (Mill.) Druce | Májovský et al. 1970 | 2n = 18 | P. multiflorum (L.) All. | Rice et al. 2014 | 2n = 28 | Polygonatum × hybridum Brügger | Asparagaceae | 130 |
| 90 | Aquatic | natural | Rice et al. 2014 | 2n = 52 | P. wrightii Morong | Májovský et al. 1987 | 2n = 52 | P. perfoliatus L | Takusagaw 1961 | 2n = 52 | Potamogeton × anguillanus Koidz | Potamogetonaceae | 131 |
| 90 | Aquatic | natural | Májovský et al. 2000 | 2n = 26 | P. pusillus | Takusagawa 1961 | 2n = 26 | P. oxyphyllus | Takusagawa 1961 | 2n = 21 | Potamogeton × apertus Miki | Potamogetonaceae | 132 |
| 90 | Aquatic | natural | Kaplan et al. 2023 | 2n =52 | P. gramineus L. | Kaplan et al. 2023 | 2n =28 | P. coloratus | Kaplan et al. 2023 | 2n =40 | *Potamogeton × billupsii Fryer* | Potamogetonaceae | 133 |
| 90 | Aquatic | natural | Rice et al. 2014 | 2n = 52 | P. wrightii Morong | Takusagawa 1961 | 2n = 52 | P. maackianus A. Benn. | Takusagawa 1961 | 2n = 52 | Potamogeton × biwaensis Miki | Potamogetonaceae | 134 |
| 90 | Aquatic | natural | Májovský et al. 1976 | 2n = 52 | P. natans L. | Májovský et al. 1978 | 2n = 52 | P. lucens L. | Kuleszanka 1934 | 2n = 52 | Potamogeton × fluitans Roth | Potamogetonaceae | 135 |
| 90 | Aquatic | natural | Ficini et al. 1980 | 2n = 26 | P. polygonifolius Pourr | Májovský et al. 1976 | 4n = 52 | P. natans L. | Preston et al. 1998 | 3n = c.39 | Potamogeton × gessnacensis G. Fisch | Potamogetonaceae | 136 |
| 90 | Aquatic | natural | Takusagawa 1961 | 2n = 26 | P. oxyphyllus Miq | Uchiyama 1989 | 2n = 28 | P. octandrus Poir | Harada 1942 | 3n = 42 | Potamogeton × kamogawaensis Miki. | Potamogetonaceae | 137 |
| 90 | Aquatic | natural | Takusagawa 1961 | 2n = 26 | P. oxyphyllus Miq | Uchiyama 1989 | 2n = 28 | P. octandrus Poir | Takusagawa 1961 | 2n = 27 | Potamogeton × kamogawaensis Miki. | Potamogetonaceae | 138 |
| 90 | Aquatic | natural | Takusagawa 1961 | 2n = 26 | P. oxyphyllus Miq | Takusagawa 1961 | 2n = 52 | P. maackianus A. Benn. | Nakata et al. 1998 | 2n = 42 | Potamogeton × kyushuensis Kadono and Wiegleb | Potamogetonaceae | 139 |
| 90 | Aquatic | natural | Kaplan et al. 2023 | 2n =52 | P. nodosus Poir. | Kaplan et al. 2023 | 2n =52 | P. gramineus L. | Kaplan et al. 2023 | 2n =52 | Potamogeton × lanceolatifolius (Tiselius) C.D. Preston | Potamogetonaceae | 140 |
| 90 | Aquatic | natural | Májovský et al. 1987 | 2n = 52 | P. perfoliatus L | Takusagawa 1961 | 2n = 52 | P. maackianus A. Benn. | Takusagawa 1961 | 2n = 52 | Potamogeton × leptocephalus Koidz. | Potamogetonaceae | 141 |
| 90 | Aquatic | natural | Májovský et al. 1987 | 2n = 52 | P. perfoliatus L | Palmgren 1939 | 2n = 52 | P. gramineus L. | Takusagawa 1961 | 2n = 52 | Potamogeton × nitens Weber | Potamogetonaceae | 142 |
| 90 | Aquatic | natural | Takusagawa 1961 | 2n = 26 | P. oxyphyllus Miq | Murín 1992 | 2n = 26 | P. berchtoldii Fieber | Nakata et al. 1998 | 2n = 28 | Potamogeton × orientalis Hagstr. | Potamogetonaceae | 143 |
| 90 | Aquatic | natural | Májovský et al. 1987 | 2n = 52 | P. perfoliatus L | Májovský et al. 1978 | 2n = 52 | P. lucens L. | Hollingsworth et al. 1998 | 2n = c.52 | Potamogeton × salicifolius Wolfg | Potamogetonaceae | 144 |
| 90 | Aquatic | natural | Kaplan et al. 2023 | 2n =52 | P. richardsonii A. Benn. | Kaplan et al. 2023 | 2n =52 | P. perfoliatus L. | Rice et al. 2014 | 3n =78 | Potamogeton ×absconditus Z. Kaplan, Fehrer et Hellq | Potamogetonaceae | 145 |
| 90 | Aquatic | natural | Kaplan et al. 2023 | 2n =52 | P. lucens L. | Kaplan et al. 2023 | 2n =52 | P. gramineus L. | Kaplan et al. 2023 | 2n =52 | Potamogeton ×angustifolius J. Presl | Potamogetonaceae | 146 |
| 90 | Aquatic | natural | Kaplan et al. 2023 | 2n =52 | P. perfoliatus | Kaplan et al. 2023 | 2n =52 | P. nodosus Poir. | Kaplan et al. 2023 | 2n =52 | Potamogeton ×assidens Z. Kaplan, Zalewska-Gałosz et M. Ronikier | Potamogetonaceae | 147 |
| 90 | Aquatic | natural | Kaplan et al. 2023 | 2n =52 | P. praelongus | Kaplan et al. 2023 | 2n =52 | P. perfoliatus L. | Kaplan et al. 2023 | 2n =52 | Potamogeton ×cognatus Asch. et Graebn. | Potamogetonaceae | 148 |
| 90 | Aquatic | natural | Kaplan et al. 2023 | 2n =52 | P. perfoliatus L. | Kaplan et al. 2023 | 2n =52 | P. crispus | Kaplan et al. 2023 | 2n =52 | Potamogeton ×cooperi (Fryer) Fryer | Potamogetonaceae | 149 |
| 90 | Aquatic | natural | Kaplan et al. 2023 | 2n =26 | P. oxyphyllus Miq | Kaplan et al. 2023 | 2n =26 | P. berchtoldii Fieber | Kaplan et al. 2023 | 2n =26 | *Potamogeton* ×*drepanoides* Z. Kaplan | Potamogetonaceae | 150 |
| 90 | Aquatic | natural | Kaplan et al. 2023 | 2n =52 | P. nodosus | Kaplan et al. 2023 | 2n =104 | *P. illinoensis* Morong | Kaplan et al. 2023 | 2n =78 | *Potamogeton* ×*faxonii* Morong | Potamogetonaceae | 151 |
| 90 | Aquatic | natural | Kaplan et al. 2023 | 2n =28 | P. polygonifolius | Kaplan et al. 2023 | 2n =52 | P. natans L. | Kaplan et al. 2023 | 2n =40 | *Potamogeton* ×*gessnacensis* G. Fisch. | Potamogetonaceae | 152 |
| 90 | Aquatic | natural | Kaplan et al. 2023 | 2n =52 | P. richardsonii A. Benn. | Kaplan et al. 2023 | 2n =52 | P. gramineus L. | Kaplan et al. 2023 | 2n =52 | Potamogeton ×hagstroemii A. Benn. | Potamogetonaceae | 153 |
| 90 | Aquatic | natural | Kaplan et al. 2023 | 2n =28 | P. zosteriformis | Kaplan et al. 2023 | 2n =26 | P. strictifolius | Kaplan et al. 2023 | 2n = 27 | Potamogeton ×haynesii Hellq. et G. E. Crow | Potamogetonaceae | 154 |
| 90 | Aquatic | natural | Kaplan et al. 2023 | 2n =52 | *P. wrightii* | Kaplan et al. 2023 | 2n =52 | *P. distinctus* | Kaplan et al. 2023 | 2n =52 | *Potamogeton* ×*malainoides* Miki | Potamogetonaceae | 155 |
| 90 | Aquatic | natural | Kaplan et al. 2023 | 2n =52 | P. perfoliatus L. | Kaplan et al. 2023 | 2n =52 | P. gramineus L. | Kaplan et al. 2023 | 2n =52 | *Potamogeton* ×*nitens* Weber | Potamogetonaceae | 156 |
| 90 | Aquatic | natural | Kaplan et al. 2023 | 2n =28 | *P. zosteriformis* | Kaplan et al. 2023 | 2n =26 | P. hillii | Kaplan et al. 2023 | 2n = 27 | *Potamogeton* ×*ogdenii* Hellq. et R. L. Hilton | Potamogetonaceae | 157 |
| 90 | Aquatic | natural | Kaplan et al. 2023 | 2n =28 | *P. polygonifolius* | Kaplan et al. 2023 | 2n =26 | *P. berchtoldii* Fieber | Kaplan et al. 2023 | 2n =27 | *Potamogeton* ×*rivularis* Gillot | Potamogetonaceae | 158 |
| 90 | Aquatic | natural | Kaplan et al. 2023 | 2n =52 | P. nodosus Poir. | Kaplan et al. 2023 | 2n =52 | P. natans L. | Kaplan et al. 2023 | 2n =52 | *Potamogeton* ×*schreberi* G. Fisch. | Potamogetonaceae | 159 |
| 90 | Aquatic | natural | Kaplan et al. 2023 | 2n =104 | *P. schweinfurthii* | Kaplan et al. 2023 | 2n =52 | P. crispus | Kaplan et al. 2023 | 2n =78 | Potamogeton ×serrulifer Z. Kaplan | Potamogetonaceae | 160 |
| 90 | Aquatic | natural | Kaplan et al. 2023 | 2n =104 | P. illinoensis Morong | Kaplan et al. 2023 | 2n =52 | P. gramineus L. | Kaplan et al. 2023 | 2n =78 | *Potamogeton* ×*spathuliformis* (J. W. Robbins) Morong | Potamogetonaceae | 161 |
| 90 | Aquatic | natural | Kaplan et al. 2023 | 2n -52 | P. lucens L. | Kaplan et al. 2023 | 2n =52 | P. gramineus L. | Kaplan et al. 2023 | 2n =78 | *Potamogeton* ×*torssanderi* (Tiselius) Dörfler | Potamogetonaceae | 162 |
| 90 | Aquatic | natural | Kaplan et al. 2023 | 2n =52 | P. praelongus | Kaplan et al. 2023 | 2n =52 | P. crispus | Kaplan et al. 2023 | 2n =52 | *Potamogeton* ×*undulatus* Wolfg. | Potamogetonaceae | 163 |
| 90 | Aquatic | natural | Kaplan et al. 2023 | 2n =52 | P. natans L. | Kaplan et al. 2023 | 2n =26 | P. berchtoldii Fieber | Kaplan et al. 2023 | 2n = 39 | Potamogeton ×variifolius Thore | Potamogetonaceae | 164 |
| 90 | Aquatic | natural | Kaplan et al. 2023 | 2n =52 | P. praelongus | Kaplan et al. 2023 | 2n =52 | P. natans L. | Kaplan et al. 2023 | 2n =52 | Potamogeton ×vepsicus A. A. Bobrov et Chemeris | Potamogetonaceae | 165 |
| 90 | Aquatic | natural | Kaplan et al. 2023 | 2n =28 | P. zosteriformis | Kaplan et al. 2023 | 2n =26 | P. berchtoldii Fieber | Kaplan et al. 2023 | 2n = 27 | Potamogeton berchtoldii × P. zosteriformis | Potamogetonaceae | 166 |
| 90 | Aquatic | ?? | Kaplan et al. 2023 | 2n=52 | P. natans L. | Kaplan et al. 2023 | 2n =52 | P. distinctus | Kaplan et al. 2023 | 3n =78 | Potamogeton distinctus × P. natans | Potamogetonaceae | 167 |
| 90 | Aquatic | ?? | Kaplan et al. 2023 | 2n =26 | P. pusillus | Kaplan et al. 2023 | 2n =26 | P. friesii | Kaplan et al. 2023 | 2n =26 | Potamogeton friesii × P. pusillus | Potamogetonaceae | 168 |
| 90 | Aquatic | natural | Kaplan et al. 2023 | 2n =26 | P. friesii | Kaplan et al. 2023 | n =52 | P. crispus | Kaplan et al. 2023 | 2n =65 | Potamogeton x lintonii Fryer | Potamogetonaceae | 169 |
| 90 | Aquatic | natural | Kaplan et al. 2023 | 2n =104 | P. illinoensis Morong | Kaplan et al. 2023 | 2n =52 | P. amplifolius | Kaplan et al. 2023 | 2n =78 | Potamogeton x luxurians Z. Kaplan | Potamogetonaceae | 170 |
| 90 | Aquatic | natural | Kaplan et al. 2023 | 2n =52 | P. perfoliatus L. | Kaplan et al. 2023 | 2n =26 | P. lucens L. | Kaplan et al. 2023 | 2n =52 | Potamogeton x salicifolius Wolfg. | Potamogetonaceae | 171 |
| 86 | Geophyte | natural | Rice et al. 2014 | 2n = 18 | S. forbesii (Baker) Speta | Greilhuber and Speta 1985 | 2n = 18 | S. bifolia L. | Speta 1971 | 2n = 18 | Scilla × allenii (G. Nicholson) Speta | Asparagaceae | 172 |
| 47 | Graminoid | natural | Hoshino et al. 1993 | 2n=42 | S. triqueter (L.) Palla | Jarolímová and Hroudová 1998 | 2n=54 | Bolboschoenus planiculmis (F. Schmidt) T. V. Egorova | Fang 1992 | 2n = 64 | Scirpus × mariqueter Tang and F. T. Wang | Cyperaceae | 173 |
| 17 | Geophyte | natural | D'Emerico et al. 2000 | 2n =36 | S. parviflora Parl. | Bernardos et al. 2004 | 2n =72 | S. lingua L. | Bianco et al. 1991 | 2n =54 | Serapias × todaroi Tineo | Orchidaeceae | 174 |
| 90 | Aquatic | natural | Kaplan et al. 2023 | 2n =78 | *S. vaginata* | Kaplan et al. 2023 | 2n =78 | *S. pectinata* | Kaplan et al. 2023 | 2n =78 | Stuckenia ×bottnica (Hagstr.) Holub | Potamogetonaceae | 175 |
| 7 | Aquatic | natural | Kaplan et al. 2023 | 2n =78 | *S. vaginata* | Kaplan et al. 2023 | 2n =78 | *S. filiformis* | Kaplan et al. 2023 | 2n =78 | *Stuckenia* ×*fennica* (Hagstr.) Holub | Potamogetonaceae | 176 |
| 7 | Aquatic | natural | Kaplan et al. 2023 | 2n =78 | S. pectinata | Kaplan et al. 2023 | 2n =78 | S. filiformis | Kaplan et al. 2023 | 2n=78 | *Stuckenia* ×*suecica* (K. Richt.) Holub | Potamogetonaceae | 177 |
| 120 | Geophyte | natural | Fedorov 1969 | 2n =26 | T. longifolia J. R. Forst. and G. Forst. | Rice et al. 2014 | 2n =66 | T. pulchella Hook. | Rice et al. 2014 | 2n =46 | Thelymitra × dentata L. B. Moore | Orchidaeceae | 178 |
| 52 | Geophyte | natural | Wang 1989 | 2n = 20 | T. tschonoskii | Grif et al. 1985 | 2n = 10 | T. camschatcense | Punina 2005 | 2n = 15 | Tillium × hagae Miyabe and Tatewaki | Melanthiaceae | 179 |
| 52 | Geophyte | natural | Wang 1989 | 2n = 20 | T. tschonoskii | Uchino and Kanazawa 1988 | 2n = 20 | T. apetalon | Haga et al. 1974 | 2n = 20 | Trillium × miyabeanum Tatew. ex J.Samej. | Melanthiaceae | 180 |
| 52 | Geophyte | natural | Grif et al. 1985 | 2n = 10 | T. camschatcense | Uchino and Kanazawa 1988 | 2n = 20 | T. apetalon | Fukuda and Horii 2004 | 2n = 15 | Trillium × yezoense Tatew. ex J. Samej. | Melanthiaceae | 181 |
| 16 | Geophyte | artificial | Lenz 1970 | 2n = 28 | T. peduncularis Lindl. | Fedrov 1969 | 2n = 32 | T. laxa Benth. | Lenz 1970 | 2n = 28 | Triteleia × tubergenii L. W. Lenz | Asparagaceae | 182 |
| 40 | Aquatic | natural | Májovský et al. 1974 | 2n = 30 | T. latifolia L. | Májovský et al. 1976 | 2n = 30 | T. angustifolia L. | Smith et al. 1967 | 2n = 30 | Typha × glauca Godr | Typhaceae | 183 |
| 103 | Graminoid | artificial | Pinto de Paula 2017 | 4n =36 | *U. decumbens* | Pinto de Paula 2017 | *2n =18* | *U. ruziziensis* | Pinto et al. 2017 | 4n =36 | *U. ruziziensis* × *U. decumbens* | Poaceae | 184 |
| 103 | Graminoid | artificial | Pinto de Paula 2017 | 4n =36 | *U. brizantha* | Pinto de Paula 2017 | *2n =18* | *U. ruziziensis* | Pinto et al. 2017 | 4n =36 | *Urochloa ruziziensis* × *U. brizantha* | Poaceae | 185 |
| 89 | Epiphyte | artificial | Hartati et al. 2024 | 2n =40 | V. dearei Rchb.f. | Hartati et al. 2024 | 2n =38 | *V. celebica* Rolfe | Hartati et al. 2024 | 2n =38 | Vanda celibica x V. deari | Orchidaeceae | 186 |
| NA | Graminoid | natural | Darlington and Wylie 1955 | 2n=28 | P. monspeliensis (L.) Desf. | Májovský et al. 1978 | 2n=28 | A. stolonifera L. | Rice et al. 2014 | 2n=28 | x Agropogon lutosus (Poir.) P.Fourn. | Poaceae | 187 |
| 1 | Epihpyte | artificial | Marchant 1967 | 2n = 50 | B. nutans H. Wendl. x Regel | Marchant 1967 | 2n = 54 | C. beuckeri E. Morr. | Marchant 1967 | 2n = 42 | x Cryptbergia meadii | Bromeliaceae | 188 |
| NA | Geophyte | natural | Anghelescui et al. 2021 | 2n =42 | *P. albida* (L.) Á.Löve and D.Löve | Anghelescui et al. 2021 | 2n =40 | D. fuchsii (Druce) Soó | Anghelescui et al. 2021 | 2n =40 | x Pseudorhiza nieschalkii (Senghas) P.F.Hunt nothosubsp. siculorum H.Kert ́esz and N.Anghelescu | Orchidaeceae | 189 |
| NA | Geophyte | natural | Anghelescui et al. 2021 | 2n =42 | *P. albida* (L.) Á.Löve and D.Löve | Anghelescui et al. 2021 | 2n =40 | Dactylorhiza fuchsii (Druce) Soó | Anghelescui et al. 2021 | 2n =40 | x Pseudorhiza nieschalkii (Senghas) P.F.Hunt nothosubsp. siculorum H.Kert ́esz and N.Anghelescu | Orchidaceae | 190 |
| 192 | Geophyte | natural | González et al. 1980 | 2n = 48 | Z. citrina Baker | Nandi 1973 | 2n = 38 | Z. candida (Lindl.) Herb. | Raina and Khoshoo 1972 | 2n = 48 | Zephyranthes × ajax Sprenger | Amaryllidaceae | 191 |
